# Sensorimotor confidence during explicit motor adaptation

**DOI:** 10.1101/2025.10.31.685783

**Authors:** Marissa H. Evans, Michael S. Landy

**Affiliations:** Dept. of Psychology, New York University, New York, NY, USA; Center for Neural Science, New York University, New York, NY, USA

**Keywords:** confidence, adaptation, uncertainty, metacognition, sensorimotor, reaching, proprioception, reward

## Abstract

Humans can adapt to large and sudden perturbations of sensory feedback. What multisensory and motor-execution cues are used to determine confidence in action success, and do the dynamics of confidence parallel those of ongoing sensorimotor adaptation? Participants made a slicing reach through a visual target with an unseen hand, followed by a continuous judgment of confidence in reach success. For the confidence judgment, participants adjusted the size of an arc centered on the target. Larger arcs reflected lower confidence. Points were awarded if the subsequent visual feedback was within the arc, and fewer points returned for larger arcs. This incentivized attentive reporting of confidence and minimizing feedback-target distance to maximize the score. After the confidence response, visual feedback of hand position was shown at the same distance along the reach as the target. A 20 deg rotation was applied to the feedback during the central 50 trials of a block (alternating between clockwise and counterclockwise across blocks). We used least-squares cross validation to compare four Bayesian-inference models of sensorimotor confidence using prospective cues (knowledge of motor noise and visual feedback from past performance), retrospective cues (proprioceptive measurements), or both sources of information integrated to maximize expected gain (an ideal observer) with additional parameters for learning and bias. All but one of the participants used proprioception to calculate sensorimotor confidence during motor adaptation in addition to prior information. Confidence recovered exponentially to pre-adaptation levels after the perturbation ended, but at a slower rate than motor learning.

**Author Summary:** We asked participants to reach to visible locations on a tablet, without being able to see their hands. We then showed them false feedback that had been rotated away from their actual hand position. This caused them to need to aim away from their intended reach direction on later trials. Confidence in their ability to have the feedback land close to the target was reduced for more trials than it took them to adapt to the rotation. To perform well, it was necessary for people to take into account their success on previous trials as well as where they sensed their hand to be positioned on the current reach.

## Introduction

To successfully interact with the physical world around us, it is necessary to adapt to changing conditions. When we experience a mismatch between the predicted outcome of an action and sensory feedback, humans make conscious and unconscious adjustments to the motor plan for subsequent movements to improve the chance of successfully achieving their goal. When evaluating confidence in the success of a motor action, humans depend heavily on prior experience. However, previous research has shown that humans do not always take action-specific information, such as proprioception, into account when making a sensorimotor confidence judgment. This is especially common if they have high proprioceptive uncertainty, although they are able to use proprioception when directly prompted (1). While action-specific information may be ignored in a static environment with fully veridical movement feedback, action-specific information becomes far more valuable when an external perturbation requires adaptation of the motor plan (2, 3). In this work we examine how sensorimotor confidence is affected by motor adaptation and what sensory information is taken into account when reacting to an environmental shift.

### MOTOR ADAPTATION

Motor adaptation is a form of error-driven learning that occurs quickly and automatically, allowing the observer to adjust the motor plan to achieve accurate movement in response to changing environmental conditions. In error-based learning, discrepancies between the expected and actual outcome of a movement are detected and gradually reduced over successive movements (4–6). Adaptation is often spurred by perturbations, which are systematic manipulations of the scene or of the moving limb that elicit an adjustment of the motor command for subsequent movements in response.

Adaptation can be explicit ⏤ a conscious, strategic change of the motor plan ⏤ or implicit. Implicit adaptation is an automatic adjustment of motor behavior that occurs without explicit awareness or intention. Implicit motor learning suggests that the brain updates internal models of movement dynamics to improve accuracy without conscious effort. In contrast, explicit adaptation involves conscious awareness and deliberate strategies to adjust motor behavior in response to perceived errors or changes in task demands (7, 8). Explicit adaptation is more rapid and flexible compared to implicit adaptation, as it can involve cognitive processes such as attention, decision-making, and explicit error correction (9–11) and is usually responsible for larger motor shifts than implicit adaptation. Explicit adaptation to a feedback perturbation involves conscious awareness of the perturbation and a deliberate adjustment to the movement plan to compensate for it (12, 13). Feedback-guided adaptation results in ongoing adjustment of motor commands based on monitoring movement outcome, which helps to achieve stability and consistency in motor performance (14).

In this work we focus on explicit adaptation by using a large, sudden perturbation to reach endpoint feedback that is noticeable to the observer. Although they are not informed in advance of the size or direction of the perturbation, nor when it will commence, they are told that a perturbation will occur. Since implicit adaptation is often small and subconscious, we opted to focus on explicit adaptation to ensure that the participant was aware of possible increased uncertainty during adaptation and could report their resulting reduction in confidence. To further induce the use of explicit adaptation, endpoint feedback was provided after the confidence report. This introduces a delay between the movement and feedback that reduces implicit adaptation. While continuous on-line feedback has been shown to produce faster and stronger adaptation (15), delayed endpoint feedback is still able to elicit explicit motor adaptation (16).

### SENSORIMOTOR CONFIDENCE

Sensorimotor confidence refers to an individual’s subjective feeling of success when attempting an action with a sensory-directed goal (17). This concept integrates sensorimotor control, motor awareness, sensory feedback, perceptual accuracy, past experience and motivational factors to shape how individuals perceive and approach motor activity. This differs from direct motor confidence (e.g., I am confident I can do a flip) because of the external goal. Importantly, the confidence report is contingent on the success of the goal. For example, if the target was missed by a large margin, the participant may confidently know where their hand is located, but report low confidence in the success of their reach since they know the target is far from their hand.

Sensorimotor confidence plays a crucial role in motor adaptation. When individuals receive accurate and timely feedback about their performance, it can boost their confidence, which in turn enhances motor adaptation (18). This highlights the close relationship between sensorimotor confidence and the processing of sensory feedback during adaptive motor learning. Additionally, uncertainty in learning has an impact on sensorimotor confidence and subsequent adaptation. Individuals adjust their confidence level based on the uncertainty of task demands and sensory feedback, which in turn influences the degree to which they engage in adaptive motor learning (19).

Furthermore, individuals who have greater confidence in their motor ability tend to adapt more quickly to changes in task demands. This suggests that higher sensorimotor confidence may facilitate faster adjustment of the motor plan when faced with perturbation or a novel environment. Conversely, lower confidence may lead to a slower adaptation rate because one may be more hesitant to modify the motor strategy (20).

Very little research has attempted to model the interaction between sensorimotor confidence and motor adaptation. Hewitsen and colleagues (21) demonstrated that confidence in motor performance is not solely based on sensory feedback from the current trial, but is also shaped by prior experience of errors during the task. This suggests that individuals integrate past error information into their metacognitive assessment, adjusting confidence accordingly as they complete visuomotor tasks. In their task, confidence was reported before any action was performed, depending only on past experience. Here, we aim to expand upon this approach and allow for trial-specific information, such as proprioception, to influence the confidence judgment. In our task, after the reach participants make a confidence judgment about the success of the reach.

In order to query the participant’s confidence without providing visual information about movement success, endpoint feedback was only provided after the confidence report. This allows self-generated trial-specific perceptual information (e.g., proprioception) to be used in the confidence judgment. Additionally, by using endpoint feedback we can employ a continuous scale for confidence in the outcome of the movement. We incentivize the confidence report by associating a wager with the confidence judgment. This allows us to compare performance to models that attempt to maximize expected gain. In addition, the amount of adaptation scales with the level of success (22, 23), thus we are using continuous confidence-judgment and feedback-error metrics instead of a binary hit or miss to elicit changes in behavior.

### CUES TO SENSORIMOTOR CONFIDENCE

Prospective cues are anticipatory signals or information available prior to execution of a motor action. These cues provide individuals with expectations about the upcoming demands and requirements of the task (24). Information about the context in which the action will occur, such as target position, the presence of obstacles, movement constraints, or expected changes in visual feedback (e.g., perturbations) provide prospective cues that shape sensorimotor confidence. Past experience and beliefs about one’s own capabilities in similar tasks serve as prospective cues.

Prospective cues contribute to the creation of expectations and predictions about task performance, influencing initial confidence levels and readiness to engage in motor activity (25–27). Lower prospective confidence has been shown to increase reaction time (28).

Retrospective cues are available after an action has been performed and are specific to that action (29). Individuals use retrospective cues to evaluate their performance on a given action. Based on feedback or other sensory information obtained after execution of a movement, they can assess how confident they are that they were successful (30, 31). Positive outcomes (e.g., accurate movement execution that achieved a goal) generally increase confidence, while negative outcomes (e.g., errors or failures) can decrease confidence (32, 33).

Each of the signals available at these two time points have their own associated uncertainty. This includes visual signals about target location or movement endpoint, motor uncertainty in the execution of a movement, or proprioceptive uncertainty regarding the body’s location during and after the action. The corresponding variances of these signals differ across individuals, making some cues more reliable than others. Given a strong dependence on visual information, prospective cues are widely used by nearly all individuals. However, people with high proprioceptive uncertainty are less likely to take retrospective cues into account (1). In an environment in which retrospective information is more valuable, such as when adapting to a perturbation, will participants be more likely to use retrospective cues when making a sensorimotor confidence judgment? The work presented here takes both retrospective and prospective cues into account when modeling confidence judgments made in tandem with motor adaptation.

### PROPRIOCEPTION

Proprioception, the sense of the body’s location in space, encompasses sensory feedback from muscles, tendons, joints, and skin that provide information about the body’s position and movement (34). There is considerable variability of proprioceptive uncertainty across individuals. This variability of proprioceptive uncertainty across participants is greater than that of visual localization (1, 35). In most cases a single multi-sensory estimate of limb position is formed by weighing and combining visual and proprioceptive signals of limb position to minimize the overall variance (36, 37). However, by removing visual feedback we can focus directly on the influence of proprioception on sensorimotor confidence.

Proprioceptive adaptation refers to the process by which proprioception recalibrates the sensory signals related to limb position, movement, and force (38). This adaptation can occur when proprioceptive signals are integrated with visual feedback to update internal models of movement and optimize motor plans (39). This sensory feedback is crucial for motor control, allowing individuals to execute precise and coordinated movements. Similar to motor adaptation, proprioceptive adaptation is influenced by feedback mechanisms and prior experience and often relies on error-based learning mechanisms (40). When there is a discrepancy between expected and actual proprioceptive feedback (e.g., due to altered limb dynamics or external perturbation), the sensory system detects the error and initiates corrective adjustment. Over time, these adjustments lead to the recalibration of proprioceptive signals to match the current environmental and task demands (41). Continuous feedback about movement outcome and sensory discrepancy facilitates ongoing adjustment of proprioceptive signals (42). Additionally, prior motor experience and learning history influence the rate and extent of proprioceptive adaptation, highlighting the role of prior experience and predictive mechanisms in sensorimotor control. In some of the models we propose here, proprioception and its related uncertainty are important parameters when evaluating confidence during adaptation. Since proprioceptive adaptation occurs even with only terminal reach endpoint feedback (43), proprioceptive adaptation is also included in our models, with updates proportional to previous feedback errors.

### CURRENT STUDY

Here, we report an experiment and associated modeling on the contribution of sensorimotor adaptation to confidence. We find that during explicit adaptation some participants rely heavily on past experience, ignoring the noisy sense of proprioception in favor of error monitoring to make their confidence judgments. However, those participants who have more reliable proprioception are more likely to take sensed hand position into account when rating confidence after adaptation. Most observers experience a period of low confidence when adjusting to a visual perturbation that lasts longer than it takes them to accurately adapt to the perturbation.

## Methods

### ETHICS STATEMENT

The study’s experimental design and recruitment process received approval from the New York University Committee on Activities Involving Human Subjects. All participants provided informed written consent and received financial compensation for their time.

### PARTICIPANTS

Sixteen self-reported right-handed individuals were selected from the New York University student body (average age: 24 years, SD: 4.9 years, 6 males). None of the participants were familiar with the experimental design. All participants had either normal or corrected-to-normal vision, no restrictions on their right arm’s mobility, and reported no motor abnormalities. Every participant completed both the control motor-awareness task (Task 1) and the primary confidence-judgment task (Task 2).

### APPARATUS

Participants were seated at a desk facing a custom-made support frame that held a mirror and a frosted-glass screen above the table’s surface (Fig. 1). A projector (Hitachi CP-X3010) mounted above the screen facing downward displayed visual stimuli on the glass screen, which was positioned above the mirror. The mirror was placed equidistant between the screen and an input tablet device (Cintiq 22; Wacom, Vancouver, WA) lying flat on the desk. The mirror reflected the underside of the screen (showing the display) back toward the participant, creating the perceptual experience that the stimuli were on the same spatial plane as the tablet surface. Participants maintained a fixed head position, with their line of sight above the mirror but below the screen, and were unable to see their hands during the task. The stimuli were presented against a mid-gray background. Reach endpoints were recorded using a stylus, held in the right hand, on the digitizing tablet at a sampling rate of 200 Hz. The experimental software was custom-built in Matlab (Mathworks, Natick, MA, USA), utilizing the PsychToolBox extension (44–46), and ran on a Dell Optiplex 9020 PC operating on Windows 10 (Microsoft, Redmond, WA, USA). Before each session, the tablet was aligned with the on-screen visuals by using a partially transparent half-silvered mirror in place of the fully silvered mirror, allowing both the display and the hand to be visible. Participants touched the tablet with the stylus at each point in a 3×3 grid of dots across the projected screen area for calibration. The resulting data were used to calculate an affine transformation between screen and tablet coordinates for each session for every participant using least-squares estimation. The calibration data from each session were only used to convert projected stimuli and recorded reach endpoints within that same session. Additionally, participants used a Griffin PowerMate control knob with their left hand for responses during both tasks.

**Fig 1:**
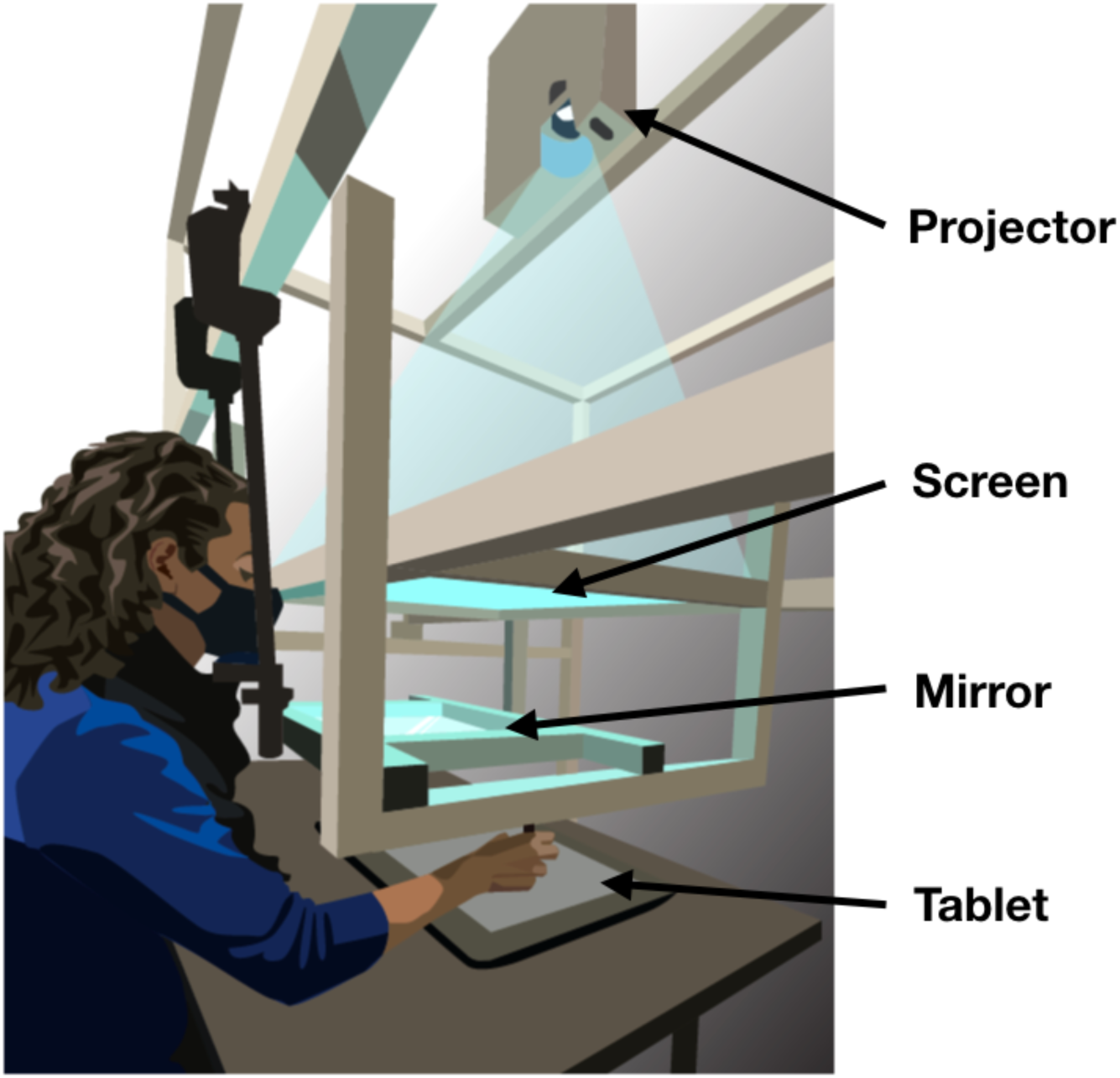
Apparatus. Participants responded with reach movements recorded by a Wacom tablet placed on a table. A mirror was suspended between a frosted glass screen and the tablet surface in the same horizontal plane. The projector above the glass screen projected stimuli downward. Participants viewed the mirror from a fixed head position and could not see the tablet or their hand while performing reaches with a stylus. The virtual image of the display was in the plane of the tablet. Figure from Fassold, Locke & Landy, (2023) Plos Comput Biol. 19(6):e1010740.

### MOTOR-AWARENESS TASK

Previous studies have demonstrated individual differences in proprioception (1, 35). The control task in our study aimed to assess the participant’s proprioceptive uncertainty separately from how it will be fit in the confidence task to provide a comparison value for each participant even if their best-fit model does not use proprioceptive error explicitly.

This task evaluated motor awareness and required participants to use proprioception, a retrospective cue, to identify reach direction. To achieve this, participants were asked to report the reach direction of unseen reaches, and a Bayesian-inference model was applied to these reports. The control task was conducted in the first session of the experiment and included twenty practice trials followed by five blocks of 60 experimental trials. Practice and experimental trials were identical to ensure consistency. However, visual feedback was provided after the judgment during practice trials and not during experimental trials.

During each trial, participants positioned the stylus within a 7 mm annulus at the bottom of the screen (Fig. 2). When the stylus was within 2 cm of the annulus center, visual feedback of the stylus location (a 4.5 mm diameter white dot) was provided. Once the cursor was inside the annulus a target (a 4.5 mm white dot) appeared 150 mm from the start location annulus, directly in front of the observer. After 800 ms, the target turned green, indicating to the participant to initiate a slicing reach through the target location. Upon completing the reach, participants were instructed to keep their right hand in place at their reach endpoint and use their left hand to indicate their perceived reach direction by rotating a dial, which moved a visual dot along an invisible arc, centered on the start point, intersecting the target. The target location remained constant throughout both practice and experimental trials. Trials were disregarded and later repeated following a visual alert if participants (1) missed the response window (800 ms to start movement from the go cue and 1200 ms to complete the reach after reach initiation); (2) moved their hand from within the annulus before the go cue; or (3) lifted their hand off the tablet surface during the reach.

**Fig 2:**
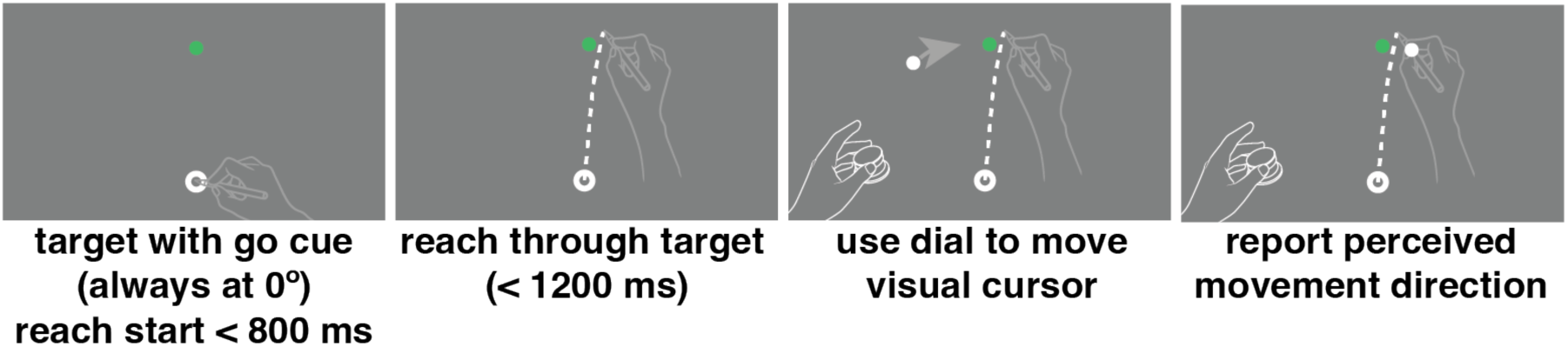
Motor-awareness task trial sequence. Participants started the trial with their hand positioned in the starting annulus at the bottom of the screen. Once the target turned green they performed a slicing reach through the target with no visual feedback of hand position. Leaving their right hand in position at the end of their reach they used the left hand to rotate a dial that moved a white dot on the screen in an arc always 150 mm from the starting point. Once the dot was positioned at their perceived reach direction they pressed down on the knob to save the report and complete the trial.

### CONFIDENCE TASK

In the primary confidence-judgment task, participants executed a slicing reach toward a visually indicated target and then rated their confidence in the accuracy of their reach using a continuous scale. Subsequently, feedback on the movement direction was provided. This approach deviates from the conventional two-alternative, forced-choice task by employing continuous measurements of motor error and a continuous confidence scale, offering a more nuanced evaluation of confidence. We employed Bayesian inference models to analyze the data and determine which parameters influence confidence assessment. Sessions 2–4 of the experiment were dedicated to this confidence-judgment task.

During a single session, participants completed a total of 400 experimental trials, divided into four blocks of 100 trials each. Prior to the experimental trials in each session, a block of 20 practice trials was conducted to allow participants to acclimate to the setup, become familiar with their own motor error, and understand the task structure. These practice trials were identical to the experimental trials but were excluded from the analysis.

On each trial, participants began by moving the stylus inside the annulus as in the motor-awareness task, this time positioned at the center of the tablet (Fig. 3). A target (a 4.5 mm green dot) appeared at a randomly chosen position on a circle (radius: 150 mm) centered on the starting point. Each quadrant of the tablet was queried an equal number of times during each block, and each direction within the quadrant was chosen uniformly and randomly. The target turned green, which served as the go cue to initiate a slicing reach through the target location as accurately as possible. For the confidence report, participants adjusted the size of an arc (radius: 150 mm from the starting annulus) using a control knob. The center of the arc itself was coincident with the target location. When adjusted, the arc’s length changed uniformly in both directions away from the target. The initial arc’s subtense was randomly selected on each trial from a uniform distribution ranging from 1 to 45 deg. Participants received points if their confidence arc enclosed the subsequently presented visual-feedback location. If the confidence arc did not include the feedback direction, zero points were awarded on that trial. If the arc did include the feedback direction, the points associated with an arc of that size were awarded. The reward was higher for smaller arc sizes, decreasing linearly from 10 points for arcs of size 2 deg or less to 0 points for 46 deg or more. Participants were required to hold the stylus at the endpoint of their reach while making the confidence judgment to encourage the use of ongoing proprioceptive signals.

**Fig 3:**
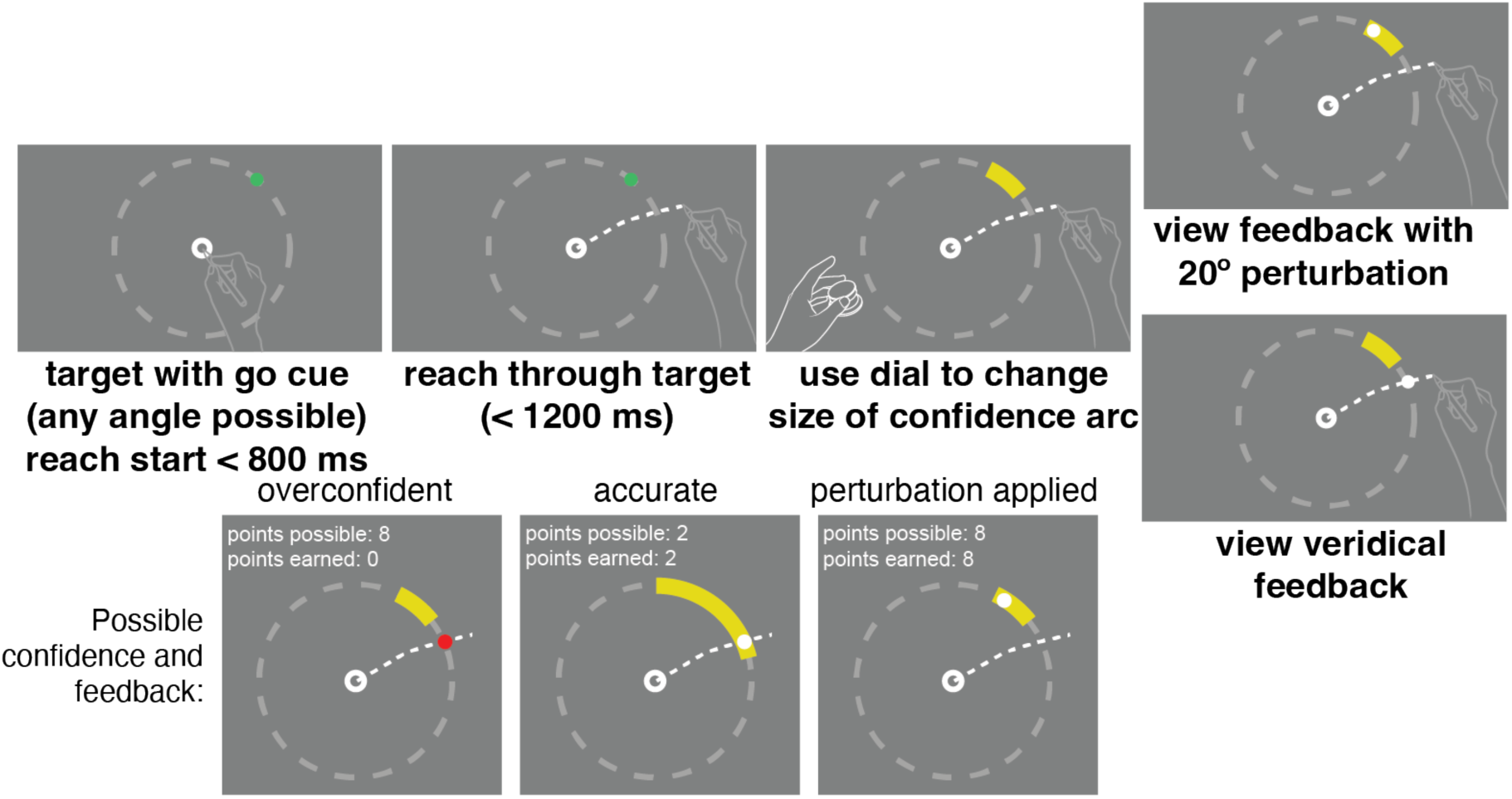
Confidence-task trial sequence and three example outcomes. Participants started with the pen positioned in an annulus at the center of the screen. At the go cue they made a slicing reach through the target without any visual feedback. Leaving their right hand at the endpoint of their reach, they used their left hand to adjust the size of a confidence arc using the control knob. The points awarded were associated with the size of the confidence arc, and points were only awarded if the feedback dot was captured by the arc. Once the confidence judgment was reported visual feedback was shown, either veridical or perturbed depending on the trial (top right). At the bottom of the figure some possible outcomes are shown. If the feedback was veridical but the confidence arc was not made large enough, thus showing overconfidence in their performance, no points would be earned (left). If the confidence arc captured the feedback location successfully (center and right) points were earned, shown here without the perturbation applied (center) and with the perturbation (right).

Trials were aborted if the participant missed the response window, moved the hand from the start location before the go cue, or lifted the hand off the tablet during the reach. In such cases, the target quadrant was shuffled back into future trials to ensure equal testing of all quadrants. No online visual feedback was provided during the reach.

A fixed 20 deg rotational perturbation was applied during trials 20-70 within each block of 100 trials, alternating clockwise/counterclockwise across blocks to reduce adaptation carryover. Participants were verbally informed at the start of the session that a rotation would occur at some point during the block and that the points awarded were contingent on the feedback location, prompting them to adjust their aiming direction to center the feedback on the target. The magnitude and direction of the perturbation were unknown to the participants at the start of each block, but could be assessed after repeated exposures.

Participants were briefed on the relationship between feedback, arc size, and points, and instructed to earn as many points as possible within these parameters. This reward structure emphasizes accurate performance (greater reward for smaller arc size) while also fostering a reasonable confidence report based on performance in each trial (1) and has been previously successful in measuring spatial-memory confidence (47).

This approach aligns the confidence report with the error scale, allowing for direct comparison between error magnitude and confidence. At the end of each block, participants were shown their running score, and at the end of the session, a leaderboard displayed all previous participants’ single-session performances, motivating participants to maximize their expected gain.

When processing the data we replaced feedback outliers where the reach was over 100° away from the target with the average of that same trial number from the other 11 runs (one trial for participant 1). We replaced outlier confidence reports greater than 45° with 45° as this is the magnitude at which the points returned dropped to zero and would return the same expected gain (one trial for participant 1). Since these outliers occurred on only one trial each we consider them to be report errors that are not indicative of the adaptation itself or the intention of the participant.

### MODELS

#### Motor-awareness task

The motor-awareness task is nearly identical to the one used in our previous paper on sensorimotor confidence (1) and we used the same model here to estimate the participant’s motor and proprioceptive noise. We assumed an ideal-performance model that combines prior knowledge of typical reach directions and the noisy sensed reach direction, and computes the maximum a posteriori (MAP) estimate of the true reach direction. The participant aims at the target direction, straight ahead (i.e., 0°) in all trials. We assume that the reach is noisy and unbiased and the distribution of reach directions has motor variance, *σ*^2^, that is known to the participant. That is, the prior distribution, *p*(*e*), of the reach direction, *e*, is *e* ∼ 𝒩(0,*σ*^2^). After the reach, the participant has a noisy proprioceptive measurement, *s*, of movement direction, with proprioceptive variance, *σ_p_*, that is also known to the participant. Thus, *s* ∼ 𝒩(*e*, *σ*^2^).

The corresponding motor and proprioceptive reliabilities are *r_m_* = 1/*σ*^2^ and *r* = 1/*σ*^2^, respectively. The participant knows and tries to estimate *e*. We assume that the variance of the visual estimate of the target direction is negligible relative to motor and proprioceptive noise and thus treat target direction as known precisely. From the participant’s perspective, the likelihood function is then *p*(*s* | *e*) = *ϕ*(*s*; *e*, *σ*^2^), where *ϕ* denotes the value of a normal distribution at the sensed direction, *s*, with a mean of the true reach direction, (unknown to the participant), and variance *σ*^2^. From the participant’s perspective, the posterior distribution is *p*(*e* | *s*) ∝ *p*(*s* | *e*)*p*(*e*), and the participant estimates the endpoint location as the maximum a posteriori (MAP) estimate of reach direction, and thus indicates direction

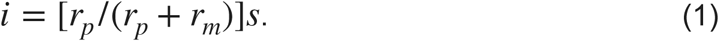

In this model motor noise, *_m_*, and proprioceptive noise, *σ_p_*, are the parameters we fit to the data. In this task we assume there is no uncertainty in reporting the perceived reach direction, that is, we do not include additional noise from using the dial to indicate the direction in the model. We estimated these parameters by maximum likelihood. Thus, from the experimenter’s perspective, the likelihood of the data, *d*, from a single trial *j*,

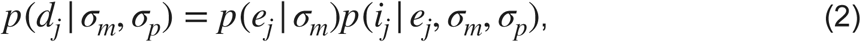

where *d_j_*= {*e_j_*, *i_j_*}, is

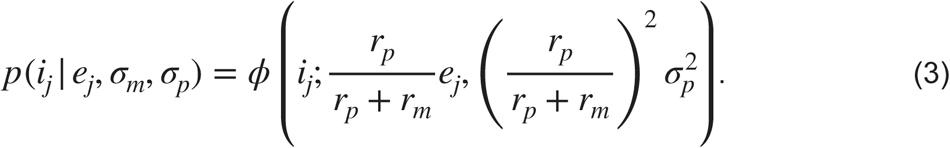

where the likelihood of the endpoint is *p*(*e_j_* | *σ_m_*) = *ϕ*(*e_j_*; 0,*σ*^2^) and the likelihood of the indicated endpoint, *i_j_*, is

Our estimates of a participant’s motor and proprioceptive noise are the values of *_m_* and *σ_p_* that maximize the overall likelihood, or equivalently the log likelihood, across all trials, {*σ_m_*, *σ_p_*^ } = arg max log *p*(*d* | *σ_m_*, *σ_p_*), where *d* is the full set of data *σ_m_*,*σ_p_* across trials. We assume independence of trials so we sum the log likelihood across all trials.

#### Confidence task

For this task we assume that the participant is explicitly aware that a perturbation may occur but does not know the size or direction of the perturbation. As a result, when updating their motor plan they must parse how much of an error was the result of their own failed reach and how much can be credited to an external perturbation applied to the feedback. Since the perturbation is quite large we assume it’s presence was known to the observer once it was introduced. Thus, a correction to the reach plan away from the target is updated trial by trial, modulated by a learning rate. Additionally the models take into account proprioception and its potential recalibration as well as a prior expectation of movement direction.

We propose three process models to compare to the data, summarized in Table 1. The output of all three is the selection of an arc size that maximizes the expected point gain for a given reach.

**Table 1.**
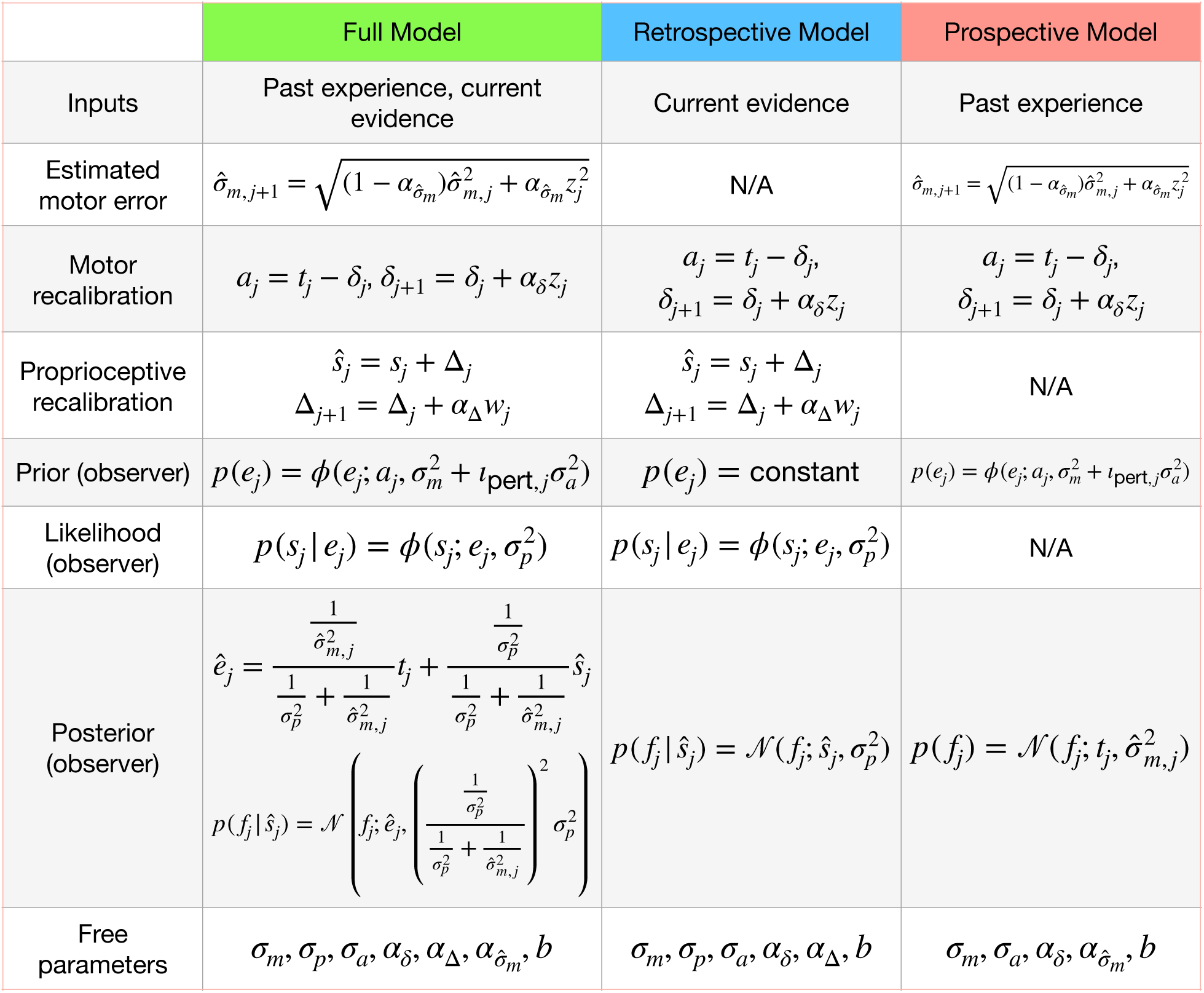
Summary of the three models.

### FULL MODEL: OPTIMAL OBSERVER MODEL WITH DELTA UPDATE

All proposed models assume that on trial *j* the participant aims in a direction, *a_j_*, that is toward the target, *t_j_*, with additional movement-plan adjustment *δ_j_* in the opposite direction of past errors:

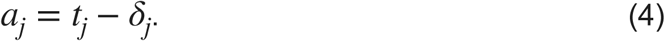

The true reach direction *e_j_* will be normally distributed around the aim point with a standard deviation that includes both motor noise, with SD *_m_*, and additional noise from uncertainty in calculation of the aim point, with SD *_a_*, only on trials after a perturbation is applied. That is, we are assuming that the perturbation is large enough to be known to the observer and results in a change in intended aiming direction that leads to additional error:

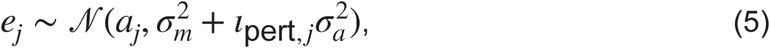

where _pert,_ *_j_* is an indicator variable that is equal to zero during the initial, unperturbed trials and one for trials following perturbed feedback. After the reach is completed, the proprioceptive measurement (i.e., the sensed reach direction) is distributed just as in the motor-awareness task:

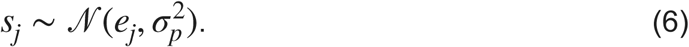

A recalibrated sensed reach direction *s_j_*^ is then calculated by shifting the proprioceptive measurement in the same direction as previous errors Δ*_j_*. Note that this adjustment is in the opposite direction to the adjustment of the aim based on previous errors (Eq. 4).

The motor plan is shifting away from the target so that the feedback lands on the target, while the sensed location needs to shift back towards the true target:

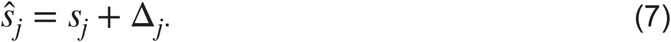

The eventual displayed feedback direction *f_j_* will be the true reach direction plus the perturbation on that trial, *p_j_*:

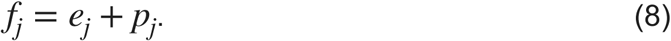

The participant’s estimate of reach direction is an average of the direction to the target, weighted by the current estimate of motor uncertainty, and the recalibrated sensed reach direction, weighted by its uncertainty:

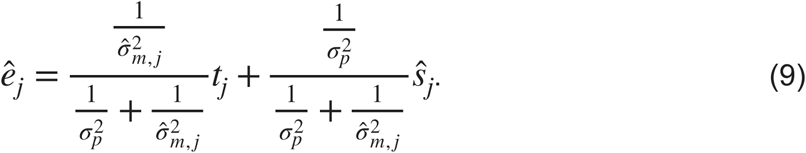

Here, the motor uncertainty is an estimate, *σ*^*_m_*, *_j_*, that incorporates recently experienced error feedback and is initiated as 0 on the first trial. This endpoint estimate and the corresponding uncertainty are used to calculate the posterior probability that the subsequent feedback will fall in any given direction:

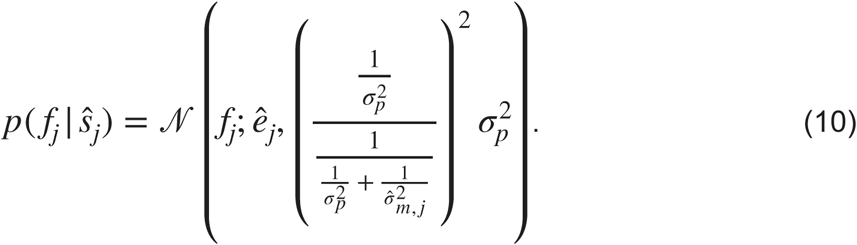

The participant selects the confidence-arc length, *l_j_*, that maximizes expected point gain. Points are awarded if the arc centered on the target with that length intersects the endpoint feedback. The number of points awarded is a linear function of arc length, reward(*l_j_*). The expected gain for arc size *l_j_* is thus:

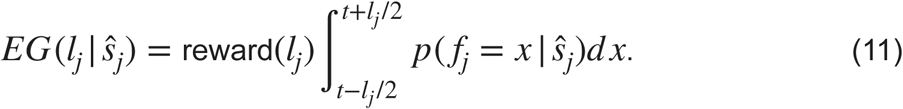

The arc length that maximizes expected gain is:

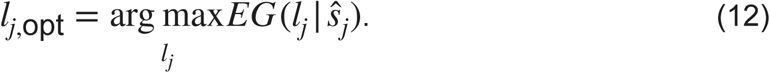

Motor adaptation to a fixed visual perturbation like ours is typically incomplete (4, 48, 49) and so we reduce the estimated feedback error by a fixed bias term during the perturbation trials. Since our data was pre-processed to align perturbation direction, is always in the coordinates that push away from ideal adaptation toward the target. This gives us *z*, the signed error between the actual endpoint feedback and the target, attenuated by a bias term *b*,

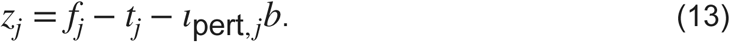

This biased error estimate is used to update the movement-plan adjustment, δ, for the subsequent trial, modulated by a learning rate *α_δ_*,

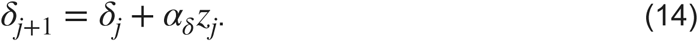

The proprioceptive recalibration is updated for the next trial in much the same way, but by using the signed error between the recalibrated sensed angle and the feedback,

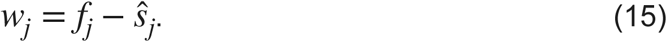

The shift applied to sensed movement direction is updated by this error, modulated by a learning rate, Δ:

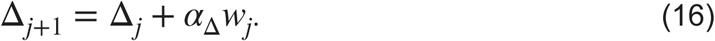

Finally, the estimate of motor error is updated to take into account the current feedback error:

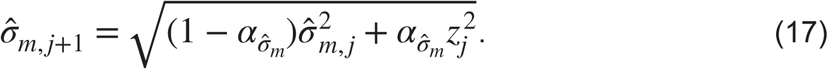

### RETROSPECTIVE CONFIDENCE MODEL

In this model reach direction and aim adaptation are still dependent on *_m_* and prior feedback, however the confidence judgment is solely dependent on the recalibrated sensed reach direction and does not incorporate knowledge of motor noise. Thus the posterior probability of the feedback is centered on the recalibrated sensed reach direction:

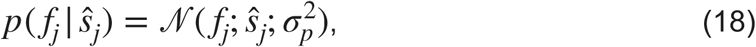

replacing Eq. 10, and Eqs. 9 and 17 are not used for this model.

### PROSPECTIVE CONFIDENCE MODEL

This model does not incorporate any proprioceptive signals so a recalibrated sensed reach direction is not calculated and the posterior probability of the feedback location is based on the estimated motor noise and the target direction alone:

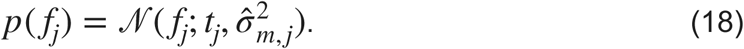

### MODEL FITS

To fit these models to the data we ran a cross-validated least-squares comparison based on 40 iterations for each model and 1000 simulations per candidate parameter set. The models were fit to the first 70 trials of each 100-trial run (twelve runs per participant), excluding the 30-trial washout period after the perturbation. We excluded the washout period as we are specifically looking at the changes to confidence during an explicit adaptation. The washout period has previously been used to investigate implicit adaptation, but is usually analyzed separately from the primary perturbation-task data (21), and requires additions to any model, as washout has different dynamics from initial adaptation.

The training data consisted of a trial-by-trial average of eleven of the twelve runs for both reach error and confidence reports. We found the best-fit parameter set based on the mean feedback error and confidence-arc sizes of the eleven runs from each participant using the comparison of an average of 1000 simulations to the data and minimizing the squared error. We then tested those parameters on the left-out run by calculating the squared error between the feedback error and confidence data from that single run and an average of 1000 simulations using the best-fit parameters chosen based on the other eleven runs. We repeated this twelve times for each model so that each run was left out one time. Given the stochastic nature of the design the comparison simulation for each parameter set was based on an average of 1000 simulations. For each of the forty iterations of the fit, we calculated the sum-of-squares value from the winning parameter set for each of the twelve runs to get a total squared error for that iteration of that model. The model with the lowest total sum-of-squares value was chosen as the winner for that iteration. The process was repeated for a total of forty iterations. The winning model for each participant was the model that won the most frequently over the forty iterations. The participant with the most winning fits within each model was selected to be the sample participant for that model in the figures shown as part of the results section.

To select the starting points for the parameters in the least-squares comparison we ran a full set of simulations across a grid of values for each of the possible parameter combinations, then compared the participant data to each of these simulations to find the parameter set in the grid that resulted in the least-squared difference. The parameter set resulting in the least-squared difference between the simulations and the data became the starting point for a BADS minimization search (50).

Using the model fits, we calculated the temporal lags in the responses to the adaptation. Specifically we calculated the first trial at which the best-fit model increased the size of the confidence arc after the perturbation was applied, the number of trials that the confidence arc size remained elevated above the pre-perturbation trials (trials 2-19) average, the percentage of confidence change from the pre-perturbation trials to during the adaptation, and at what trial the feedback error and confidence arc size stabilized after the perturbation was applied. The stabilization trial was determined by the first trial after the perturbation where the trial-to-trial difference was less than .3 in the model fits.

## Results

### MOTOR-AWARENESS TASK

The goal of the motor-awareness task was to provide estimates of each participant’s motor and proprioceptive uncertainty, *σ_m_* and *σ_p_* that was independent of the main confidence task. These estimates could then be compared to the estimates derived from the fits of the confidence models to the data from the confidence task to validate the models. This is shown in the Parameter Comparison section of our results. In the motor-awareness task participants repeatedly made ballistic reaches to a fixed target location, and then indicated the perceived endpoint direction. Parameter values for each participant were determined using maximum-likelihood estimation. Across the 16 participants the average *σ_m_* estimate was 2.98 deg (SD: 1.26 deg) and the average *σ_p_* estimate was 5.18 deg (SD: 1.97) (Fig. 4). While motor error was similar for all participants, there was a wider spread of proprioceptive uncertainty, as in our previous study (1).

**Fig 4:**
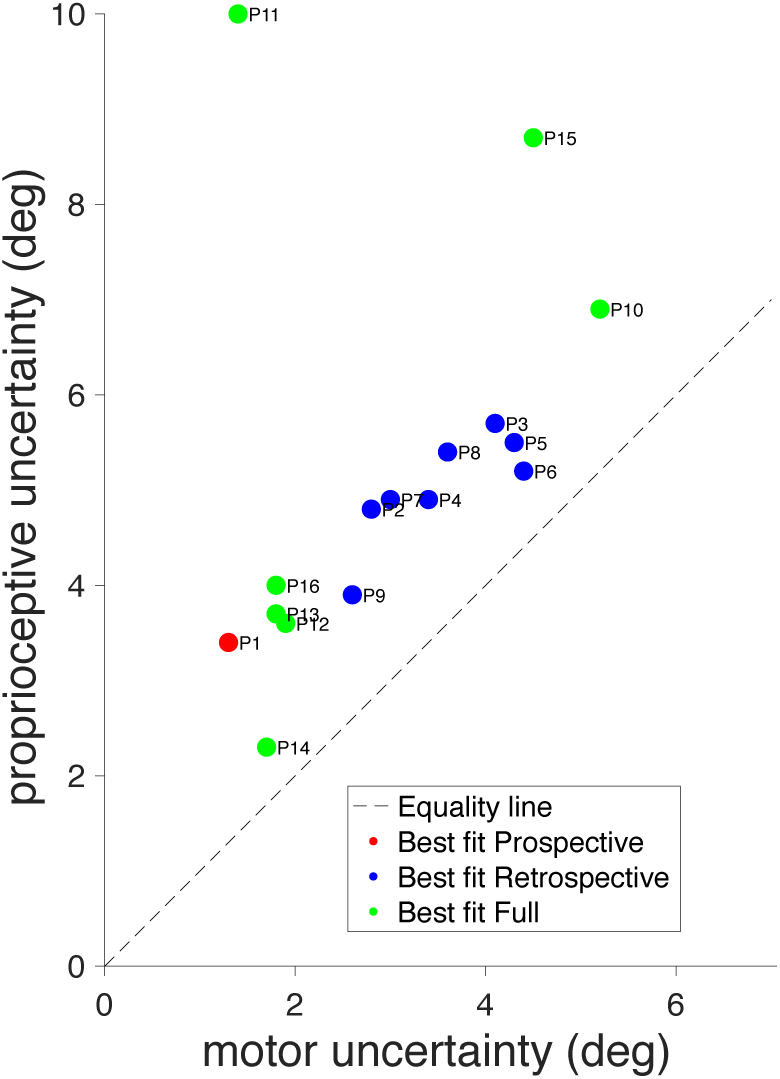
Motor-awareness task parameter comparison. Best-fit motor and proprioceptive uncertainty from the motor-awareness task for each participant. Values were fit using maximum likelihood and are colored according to which model was the best fit for that participant in the confidence task.

A similar behavioral design was used by Fassold and colleagues (1), producing parameter estimates that were found to be stable across multiple testing days. The main difference with the current study is that here a slicing style reach was used, so that reach error was one-dimensional: error in reach direction. The previous study required a pick-up-and-drop reach with the report based on a location in the 2-d plane of the tablet. All other aspects of the task remained the same and the data were analyzed in the same way.

### CONFIDENCE-TASK BEHAVIOR

The objective of our main task was to investigate how confidence judgments are affected by explicit motor adaptation and which cues are used to calculate confidence during motor learning. Participants reported confidence by adjusting the size of an arc centered on the reach target until they believed it included the direction of the subsequently presented feedback. During the middle half of each block of trials a 20° rotational perturbation was applied, either clockwise or counterclockwise depending on the block, which resulted in the displayed feedback being rotated away from the true endpoint direction. Participants were explicitly aware that a perturbation would occur but were not privy to the size, direction, or start time of the rotation prior to the session.

When the reach trajectories are split into three groups based on the trial type: veridical feedback (trials 1 to 19), perturbation (trials 20 to 70), and washout (trials 71 to 100), there is clear evidence that most participants responded to the perturbation appropriately (Fig. 5). By shifting their aiming direction, the feedback once again landed on the target after the perturbation was applied. During washout, when the perturbation was removed, the aim returned to baseline.

**Fig 5:**
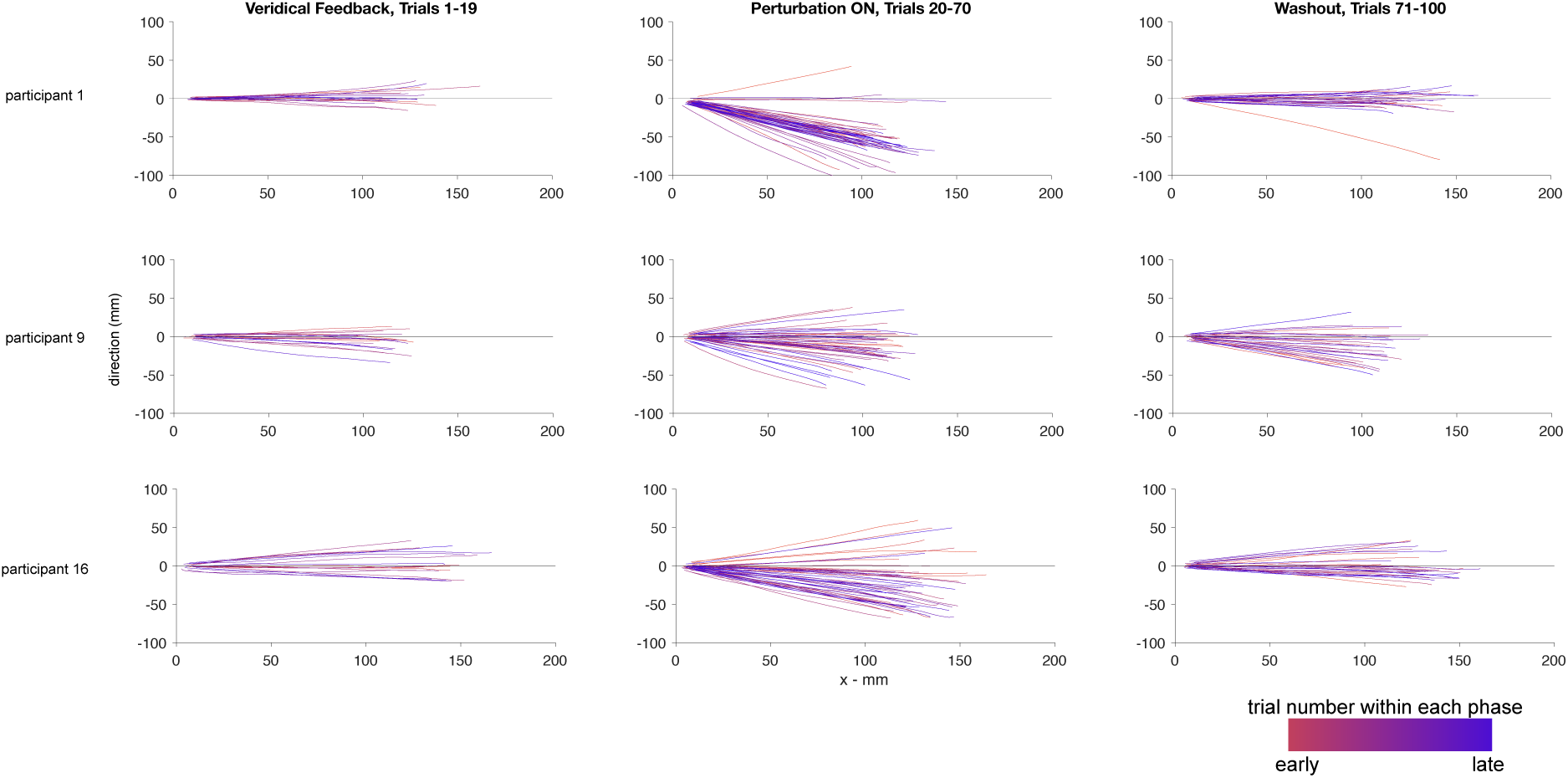
Reach trajectories from the confidence task. Reach trajectories for three sample participants from a sample block separated into the three different phases of the trial sequence, veridical feedback, perturbation on, and washout. Trajectories are rotated and flipped to align all targets and perturbation directions. Early trials within each section are shown in red while late trials are in blue.

### POINTS AWARDED

On average, participants captured 67.4% (SD: 10.4%) of the feedback locations within their confidence arc, successfully earning points on those trials. On average, participants earned 4809 points (SD: 1018) over the course of the experiment.

Theoretically perfect performance—capturing the feedback location on every trial with the smallest arc size possible—would have resulted in an average of 9662 points (SD: 662.05). Thus, participants earned 49.3% (SD: 7.5%) of the points they could have possibly earned given their reach performance.

### MODELING

We fit three models to the data using a cross-validated least-squares criterion by comparing the results of averages of stochastic simulations to each participant’s data. The three models included a prospective model that made the confidence judgment at the start of the trial and ignored trial-specific feedback, a retrospective model that did not take into account past performance but only the outcome of the current trial when calculating confidence, and a full model that used all available information to determine the confidence report. The models were fit to the first 70 trials in the block. We did not fit the data from the washout period after the perturbation was removed.

### MODEL FITTING

The full model has seven free parameters: motor uncertainty, *σ_m_*, proprioceptive uncertainty, *σ_p_*, aiming uncertainty, *σ_a_*, proprioceptive learning rate, *σ*_Δ_, motor learning rate, *α_δ_*, motor-variability learning rate, *α_σ_*_^_*_m_* and bias *b*. The retrospective model has six parameters as it did not incorporate an estimated motor uncertainty value based on previous errors, thus eliminating the motor-variability learning rate, *α_σ_*_^_*_m_*. The prospective model has five parameters as it did not use proprioception and therefore eliminated the parameters for proprioceptive uncertainty, *σ_p_*, and the learning rate for proprioceptive adaptation, *σ*_Δ_.

After running 40 cross-validated least-squares fits on each participant’s data, the model with the most wins was determined to be the best fit for each participant. One participant was best fit by the prospective model, eight by the retrospective model, and seven by the full model (Table 2). Participants 2, 3, 10 and 11 were decided by a plurality of the simulations, while the remaining participants displayed a clear majority indicating one specific winning model.

**Table 2:**
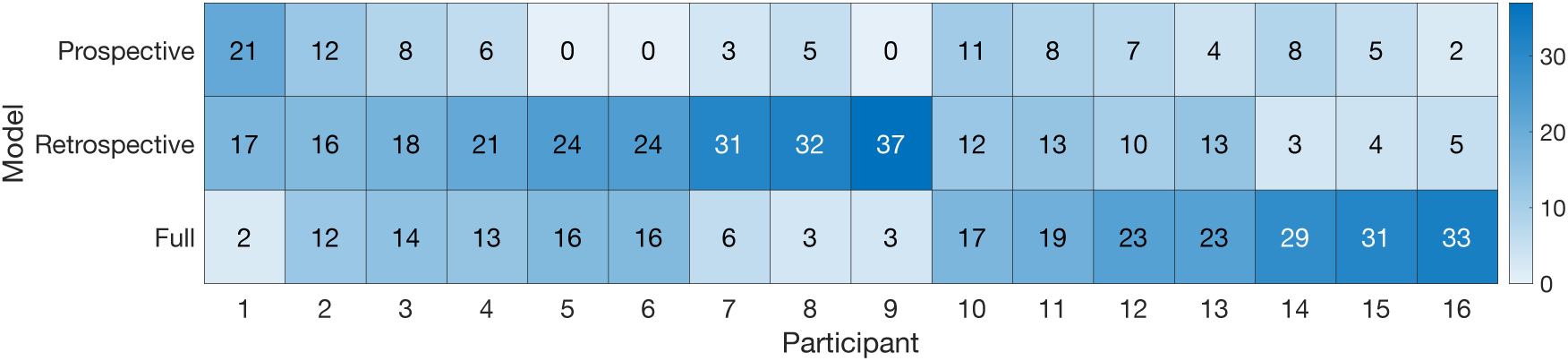
Table of winning models Forty model fits were run for each participant’s data set using cross-validated least squares and simulations. The winning model for each round was the model and parameter set that resulted in the lowest least-squares value when compared to the behavioral data. Light squares indicate fewer winning fits and dark squares signify the most fits out of the 40 possible iterations. Participant 1 was best fit by the Prospective model, 2-9 by the Retrospective model, and 10-16 by the Full model. Participants 2, 3, 10, and 11 had their winning models determined by a plurality, not a majority of the fits.

### SIMULATIONS WITH BEST-FIT PARAMETERS

To visually compare the predictions of each model to the behavioral data, we reused the results of the model fitting. For each model and participant, there were forty iterations of fitting that model to the data. For each model, we picked the parameter sets from an iteration in which that model provided a better fit than the other two models, and had the lowest total sum of squared errors of the possible winning iterations for that model. We ran a full simulation with each of the 12 parameter sets (one set for each left-out run).

The average of the 12 simulations is overlaid on top of the averaged data across the 12 runs, shown as the solid curves in Fig. 6. If a participant had zero iterations of the fit in which a given model won (Participants 5, 6 and 9; Prospective model), the plot is left blank for that participant and model combination. Shaded regions represent SEM. The grey shaded area denotes the trials where the perturbation was present. For all participants the perturbation was introduced at trial 20, resulting in a sudden decrease in the accuracy of the visual feedback, followed by an exponential decay to an asymptote (Fig. 6, blue fit curves). Plots of the individual time courses and best-fit model parameters for every participant can be found in the Supplement. These process models fit both the reach adaptation and the confidence reports well for those participants whose underlying model was a best match.

**Fig 6:**
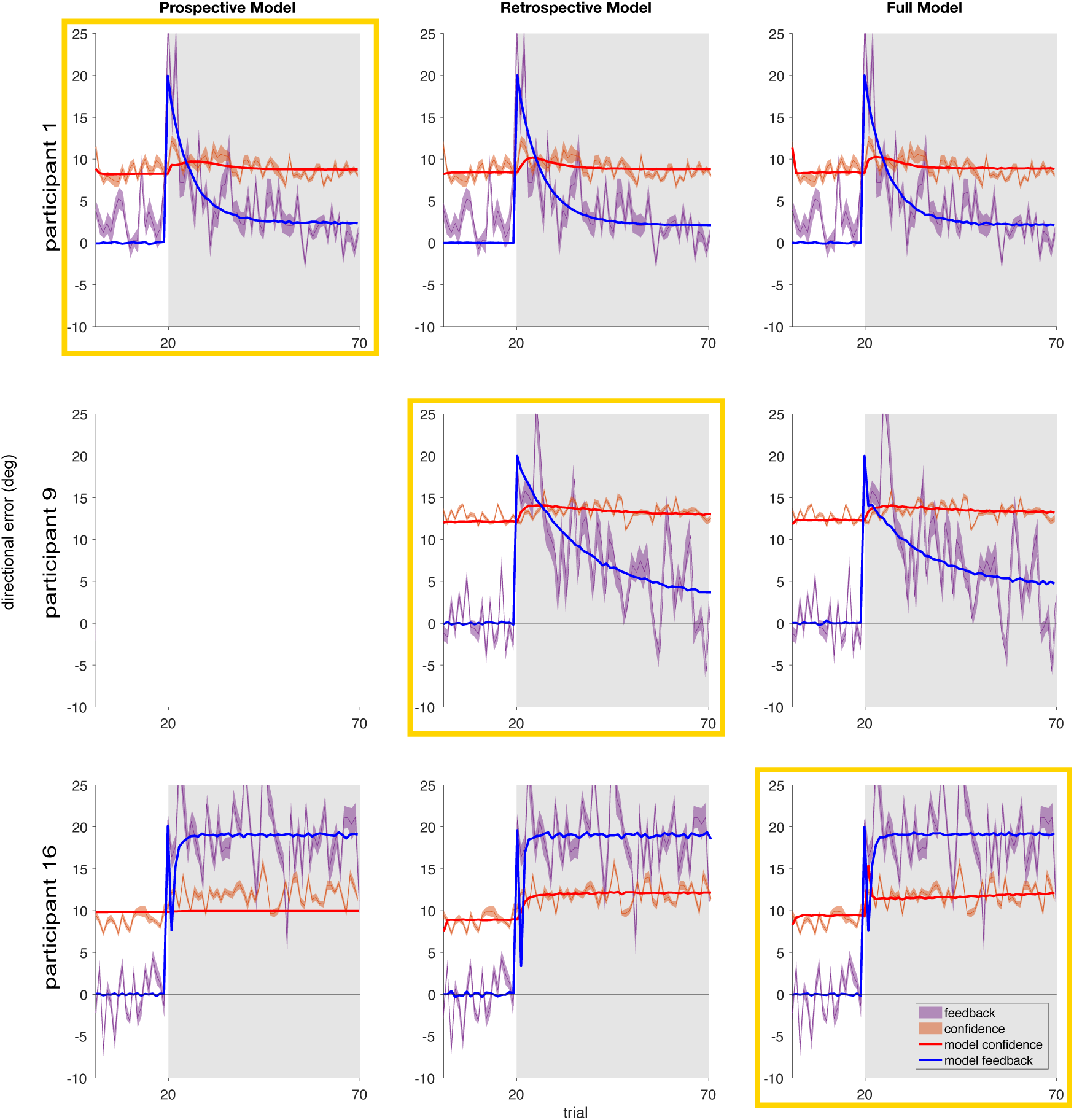
Simulations of the models using the best-fit parameters compared to the behavioral data. Three sample participants are shown. We picked one sample participant who was best fit by each model, i.e., with the greatest number of winning iterations. For each sample participant the simulated reach error and confidence report (averaged over 1000 stochastic simulations and 12 iterations) using the best-fit parameters for each model is overlaid on the behavioral data for that participant. The winning model for that participant, determined by cross-validated least squares, is highlighted with a yellow box. If a particular model never was the best fit for a given participant it has been left blank in the plots.

Taking a closer look at the fits here, explicit adaptation to the visual perturbation stabilized after an average 21.9 trials (SD: 25.8) for the Prospective model, 21.1 trials (SD: 17.1) for the Retrospective model, and 15.1 trials (SD: 13.6) for the Full model (Fig. 6, blue fit curves). Three participants fully adapted to their baseline accuracy (participants 7, 13 and 14), one participant over-corrected (participant 8), and two participants never fully stabilized their adaptation (participants 15 and 16). After the perturbation was applied we observe a lag before confidence was reduced (i.e., the confidence radius was increased). Confidence arc size (Fig. 6, red fit curves) increased progressively before stabilizing over an average of 2.4 trials (SD: 5.0) for the Prospective model, 1.8 trials (SD: 1.7) for the Retrospective model, and 2.9 trials (SD: 4.9) for the Full model after the perturbation was introduced, and in none of the models did the confidence return to the pre-perturbation level while the perturbation was applied. On average, during that period confidence arc size increased by 88.2% (SD: 9.7%) for the Prospective model, 82.2% (SD: 8.2%) for the retrospective model, and 82.5% (SD: 8.0%) for the Full model from pre-perturbation (trials 2 through 19) levels, to during the adaptation after confidence had stabilized (trials 30 through 70), representing a decrease in the participant’s sensorimotor confidence (Fig. 6, red fit curves). In general, while the rotational perturbation was applied the direction of participants’ reaches was more variable than during the trials where feedback was veridical and no adaptation was required. This suggests that additional uncertainty is added when calculating the new aim point on top of the average motor uncertainty when executing the reach prior to adaptation, which in turn lowers confidence on average.

### PARAMETER COMPARISON

The parameters estimated from the motor-awareness task (shown as black horizontal bars in the first two columns of Fig. 7), *σ_m_* and *σ_p_*, were also separately estimated for the model fits to the data from the main task. The best-fit values for these parameters were similar for most participants when they were available given the particular best-fit model (Fig. 7). This indicates that proprioceptive and motor uncertainty were well captured by our models in both tasks. Participant 9, who was best fit by the Retrospective model, was never best fit by the Prospective model, so this participant lacks a set of winning parameters for this model.

**Fig 7:**
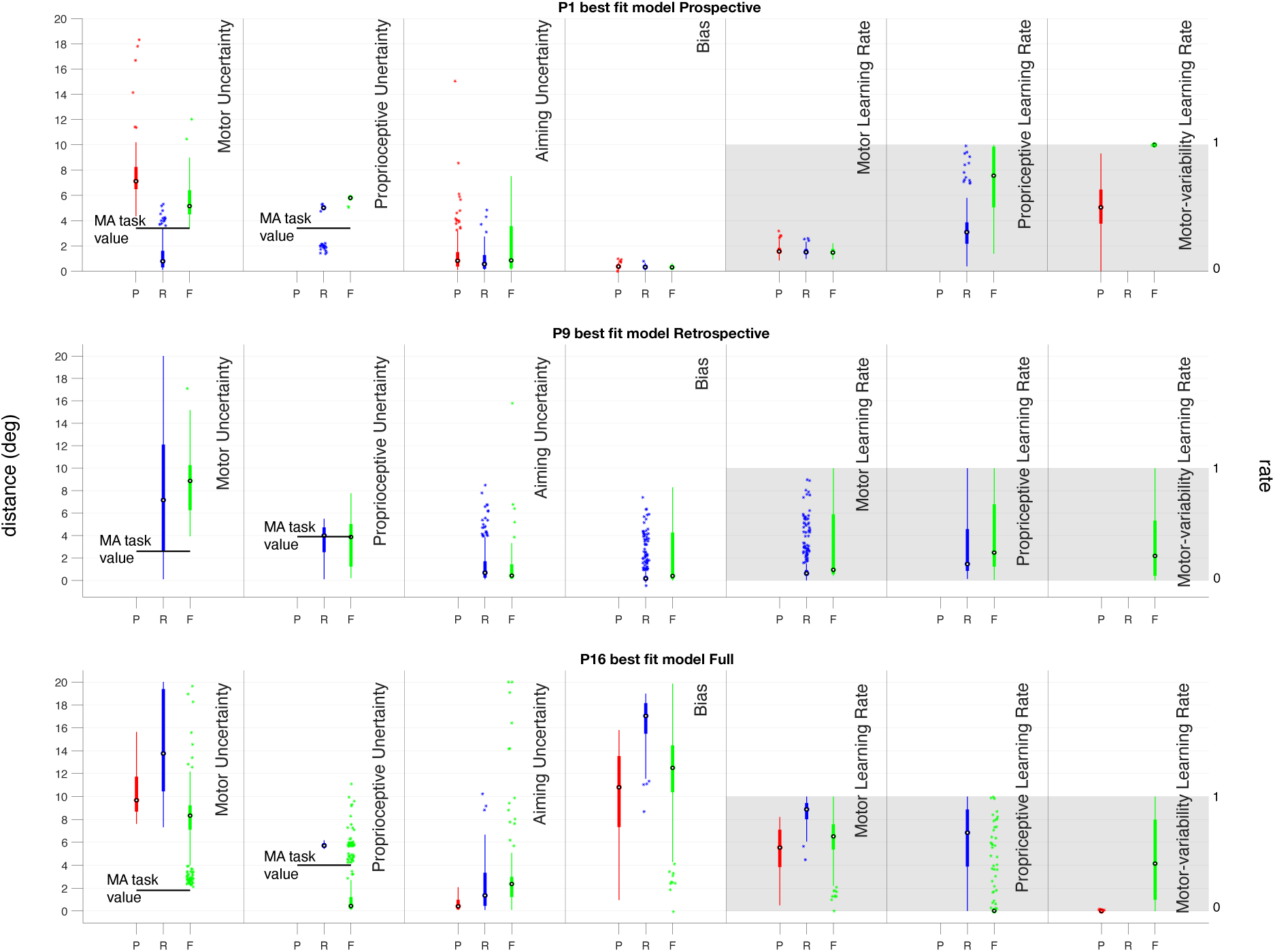
Parameter comparisons for each model fit. For each of three example participants, one best-fit by each model, we’ve plotted the range of parameters resulting in a winning iteration of a given model. On the *x*-axis the three models are represented for each parameter, Prospective (P) in red, Retrospective (R) in blue, and Full (F) in green. There are 12 sets of parameters for each iteration so there are between 24 and 444 data points for each model, depending on the number of winning iterations. These box plots show the parameters from the fits that resulted in the winning model producing the lowest least-squares value and winning that round. If the model did not use a particular parameter it has been left blank here. If that participant never had a winning iteration of a given model, there are no winning parameters to show and the section has been left blank. The box edges are the upper and lower quartiles, with the horizontal line reflecting the median, dots representing outliers (calculated using the interquartile range) and whiskers reflecting maximum and minimum values that are not outliers. The black bars in the motor- and proprioceptive-error plots are the values fit for each participant in the separate motor-awareness task. None of the data used to generate those values was used in calculating the parameters shown here and vice versa.

The parameters here explain different behavioral changes in the data. Motor error, *σ_m_*, is how noisy the participant’s reach is at baseline, when there is no additional uncertainty caused by the perturbation. Proprioceptive uncertainty, *σ_p_*, reflects how accurately they can detect their hand’s position in space, and comes into play during the confidence judgment (in applicable models) as points are awarded for the capture of the feedback location, which is a rotation of the true hand position, resulting in more accurate reports when this uncertainty is low. Aiming error, *σ_a_*, is additional uncertainty added to the reach when attempting to explicitly account for a rotational perturbation.

This accounts for the small decrease in accuracy seen in reaches made after the perturbation is applied. Bias, *b*, is the amount that the observer is drawn back toward the target after adaptation is complete, and this accounts for the plateau seen in several participants that sits a couple degrees away from perfectly counteracting the perturbation. Motor learning rate, *α_δ_*, is the rate at which previous errors are taken into account when updating the aiming direction, this parameter changes the slope of the adaptation. When the value is close to one the adaptation happens very quickly, and the lower the value is the slower the adaptation occurs. Proprioceptive learning rate, *σ*_Δ_, controls the rate at which the sensed hand location adapts toward the feedback, if this value is low the mean of the likelihood is pulled toward the true hand location and if it is high the mean of the likelihood sits near the feedback location. Given the rotational adaptation for reaches over all 360° of possible directions, this value had a large variance for all participants fits, indicating it did not make as meaningful of a contribution as the other parameters. Motor-error learning rate, *α_σ_*_m_, is the rate at which added uncertainty from the perturbation is attributed to oneself. When this value is high, more weight is given to recent mistakes, attributing more of the shift from the perturbation to one’s own error, and when this value is low the assumption of one’s own motor noise is more stable across trials. When a participant thinks they have a high motor-error uncertainty, they are more likely to report low confidence.

## Discussion

In this study we examined the process by which motor adaptation affects the calculation of sensorimotor confidence. To do so, sixteen participants performed two sensorimotor reaching tasks: a motor-awareness task to independently fit proprioceptive uncertainty, and a confidence task during a rotation-based motor adaptation. We fit three Bayesian-inference models to each participant’s data, independently for the motor-awareness task, and performed a cross-validated least-squares fit of the behavioral data to simulated data from each model for the confidence task. Idiosyncratic strategies between observers were indicated by the differences in the best-fit models. One participant was best fit by the prospective model, eight by the retrospective model, and seven by the full model. We hypothesized that during motor adaptation, participants would be more likely to use retrospective information such as proprioception, even if it was slightly unreliable. This would be shown by a higher likelihood of fit by the retrospective and full models, which is what we see here for our participants.

The variation seen between participants for their best-fit model could be driven by their underlying proprioceptive uncertainty, resulting in a lower reliability for retrospective information for some participants, leading to the additional use of prior information as we see in our Full model. There could be other sources for these differences as well, such as differences in cognitive strategy or difficulty adapting to the perturbation. There are four participants (2, 3, 10, and 11) who are best fit by a model by a plurality, not a majority, of simulated fits. This may result from additional uncertainty derived from one of these sources, or a change in strategy over sessions resulting in varied best-fit model scenarios.

The adaptation used here was explicit with all participants receiving prior knowledge about the visual rotation occurring in the feedback. While aiming error is a parameter accounted for in our model, we did not explicitly ask participants to report an aim direction prospectively. Rather, we estimate it based on their motor uncertainty and true reach direction. Because of this explicit adaptation we assume that there is negligible influence of a sense of agency on our results. Participants are aware that a rotation will be applied to the visual feedback during the task and should not have the impression that they are suddenly performing very poorly at reaching. Additionally it was very possible that participants would not augment their confidence ratings during an adaptation. They have explicit knowledge of the perturbation, additional motor noise or proprioceptive noise has not been introduced so the action itself is the same physical difficulty as the reaches performed prior to the perturbation.

We found a greater use of proprioception during confidence judgments with our motor-adaptation task than in previous research (1), but it is also important to note that this task used a slightly different behavioral paradigm that may improve access to proprioceptive signals when compared with previous work. In this task participants kept their hand at the reach endpoint position when reporting their confidence using their non-dominant hand, instead of returning to the starting point prior to the report. Having the online proprioceptive signal available during the confidence judgment may allow it to be more prominently used by participants who are likely to use it anyway.

Previous research has shown that adaptation is more prevalent with full reach-trajectory feedback (4, 15, 16, 43, 49, 51, 52). However, since the confidence judgment relied on estimation of the endpoint location, providing feedback prior to the confidence judgment was not possible. Given that only endpoint feedback was provided, and delayed after reach termination, it could take additional cognitive effort to couple the feedback with the reach, reducing the amount of adaptation that was possible.

Decoupling the feedback from the true reach trajectory, such as in an error-clamp paradigm (4), may be an interesting way to investigate the implicit effects of adaption on confidence and would be worth researching further. The washout period of the trial block, after the perturbation has been turned off, has been previously investigated as a cue to implicit adaptation (4, 53). In the current study we did not model the washout trials as we were specifically looking at explicit adaptation, however there may still be additional processes at work implicitly.

To earn points in this paradigm the participant needs to capture the perturbed feedback location within the confidence arc. This means the participant needs to have high confidence in their motor awareness, and also their explicit representation of the size and direction of the perturbation. This differs from work where the aiming direction is reported with visual feedback prior to the reach commencing, thus removing motor awareness as a possibly influence on the report and associated confidence (4, 5, 11, 21). In these cases only prospective sources of uncertainty influence confidence. It’s possible that having the time to perfect the aiming direction before the reach would reduce some of the decreased confidence seen when the perturbation was in effect.

The fast nature of the trial’s time requirements inherently adds uncertainty, when additional aiming calculations are needed quickly to account for the rotation, greater errors can be expected. The aiming-error parameter *σ_a_* in all models accounts for a miscalculation of the perturbation magnitude, and the feedback-learning, *σ*_Δ_ in both the Prospective and Full models, accounts for the amount that a participant might change their assumption of the magnitude of the perturbation, and thus their aiming direction based on prior feedback. Additionally some participants showed an adaptation bias, where their adaptation magnitude plateaued at a given distance away from optimal. There may be additional influences on sensorimotor confidence not included in our models. However, given what is measurable with our apparatus and behavioral paradigm, the process models presented here do well at replicating the effects seen in our data. The estimate of the learning rate for proprioceptive uncertainty is highly variable for all the models in which it was used (Retrospective and Full). The adaptation task used targets located across all 360° of directions, making adaptation difficult across such a large workspace. It’s likely that while explicit adaptation in rotation and gain can be universally extrapolated (54, 55), proprioceptive adaptation may not follow the same universal coordinate shifts. We tested target locations in all directions from the central starting point.

Therefore, across trials the participant was reaching to the left, to the right, away or towards themselves. As the perturbation was a rotation, the error direction flipped when reaching toward oneself in the bottom half of the possible target locations, versus when reaching forward. Accounting for this flip mentally when generating a new aim point can be difficult, and may be responsible for the increased error seen during the perturbation trials. Additionally natural biomechanics show improved strength and stability when reaching toward the body versus away (56–58), which may also have some effect on reach accuracy. Prior to centering the data during pre-processing we checked to make sure there was no increase of average error for a particular spatial direction (each direction considered as a 45 degree increment of possible target directions) over another during the perturbation, so the increase in uncertainty is likely tied to the rotation itself and not a specific reaching direction. Therefore, for our analyses we rotated all trials target locations into a single direction, and flipped the direction of half the trials so all perturbations were facing the same direction for the analysis.

From the perspective of the participant, it’s possible that the increased difficulty in the task resulted in a reduction of confidence. Reaching straight toward a target is much easier than making a split second decision when a target appears to reach either to the left or right of it depending on where it falls in relationship to your hand to account for the rotation, then execute a reach without a visible target, but a mentally placed aim point instead. The amount of uncertainty they might have had before the trial started was already heightened, before adding in the motor and proprioceptive error that comes with the reach itself. Additionally, the proprioceptive mismatch between the endpoint feedback and hand position, although known to the observer, could prompt the use of larger confidence arcs as it may be difficult to ignore the proprioceptive signal from the hand entirely, especially if proprioceptive recalibration was low (59, 60).

## Conclusion

We have demonstrated that sensorimotor confidence is reduced following an explicit adaptation. Additionally individuals with lower proprioceptive uncertainty are more likely to incorporate their sensed hand position when making a confidence judgment, while others instead depend heavily on the average of past errors.

## Author Contributions

MHE carried out the formal analysis, investigation, software, data curation and visualization, and wrote the original manuscript. MHE and MSL collaboratively worked on conceptualization, methodology and editing. MSL and MHE are responsible for funding acquisition.

## Data Availability

All data and code files will be made available on OSF database.

## Funding

This work was supported by NIH grant EY08266, NIH Training Program in Computational Neuroscience (TPCN) grant T90DA059110, and the NYUAD Center for Brain and Health, funded by Tamkeen under NYU Abu Dhabi Research Institute grant CG012. The funders had no role in study design, data collection and analysis, decision to publish, or preparation of the manuscript.

## Competing Interests

The authors have declared no competing interests exist.

## Acknowledgments

These computational analyses were supported by NYU High-Performance Computing resources and services.

---

**Supplement 1:**
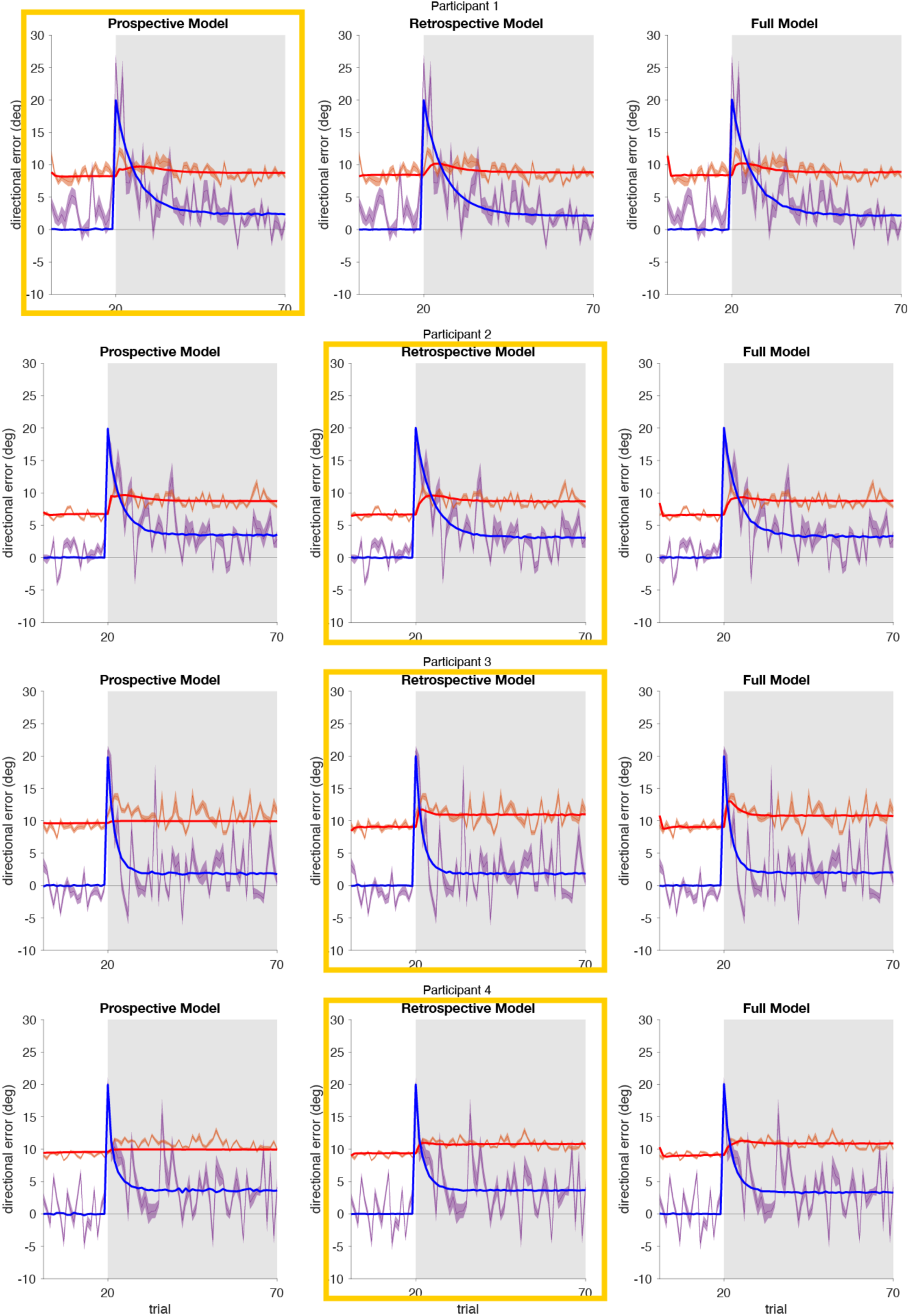
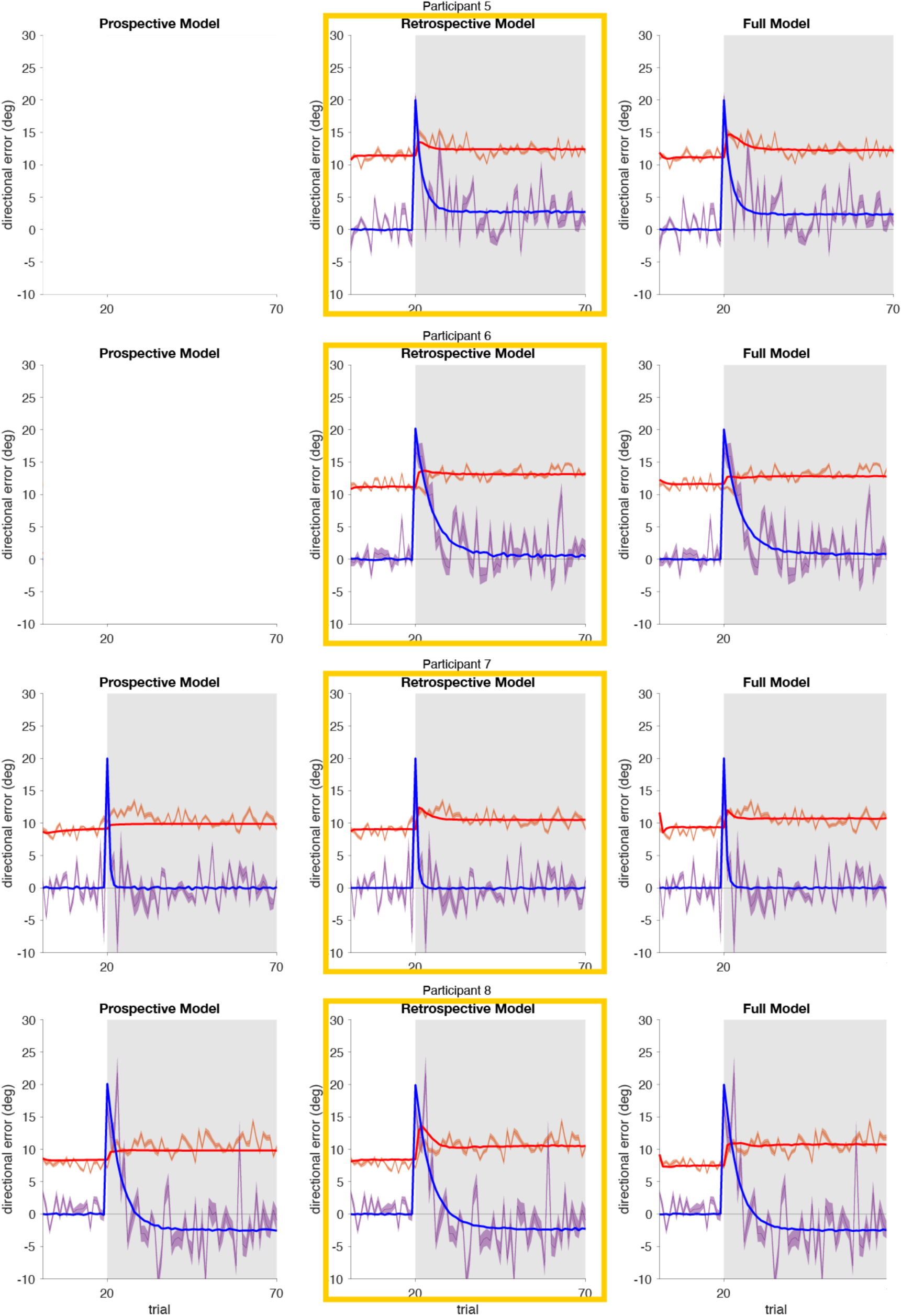
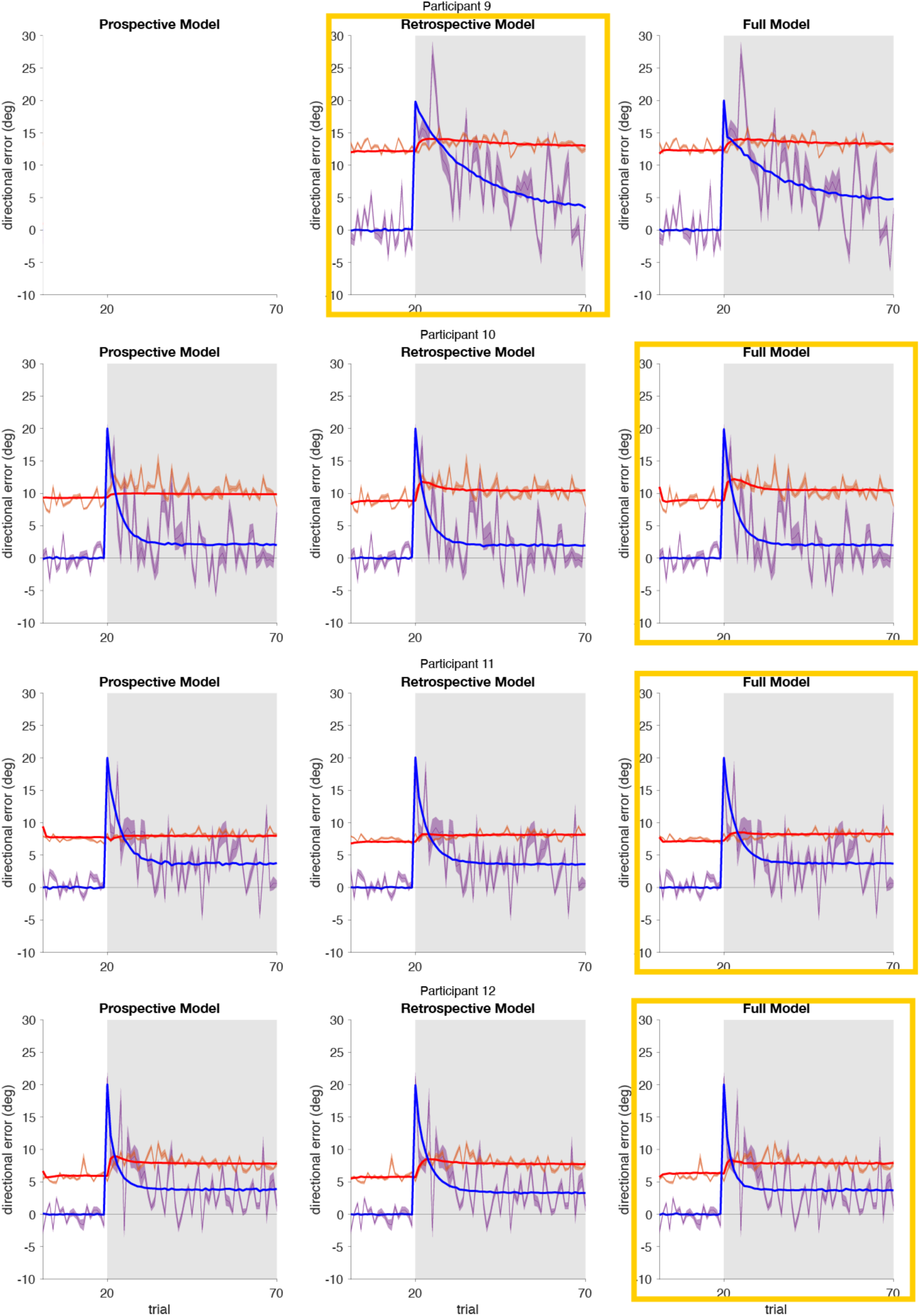
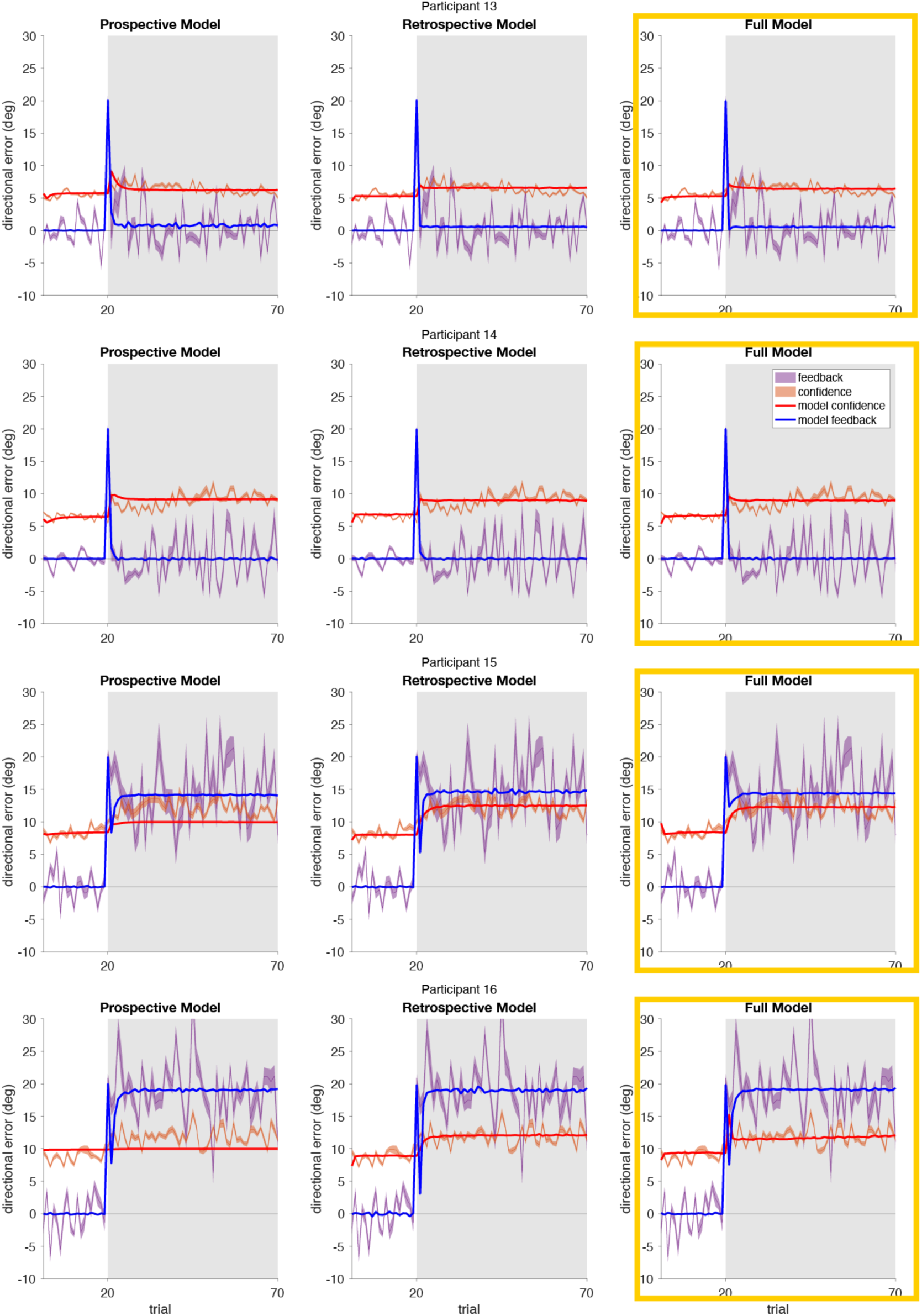
Simulations of the models using the best-fit parameters compared to the behavioral data. For each participant the simulated reach error (blue) and confidence report (averaged over 1000 stochastic simulations and 12 iterations, red) using the best-fit parameters for each model is overlaid on the behavioral data for that participant (reach error in purple, confidence in orange). The winning model for that participant, determined by cross-validated least squares, is highlighted with a yellow box. If a particular model never was the best fit for a given participant it has been left blank in the plots.

**Supplement 2:**
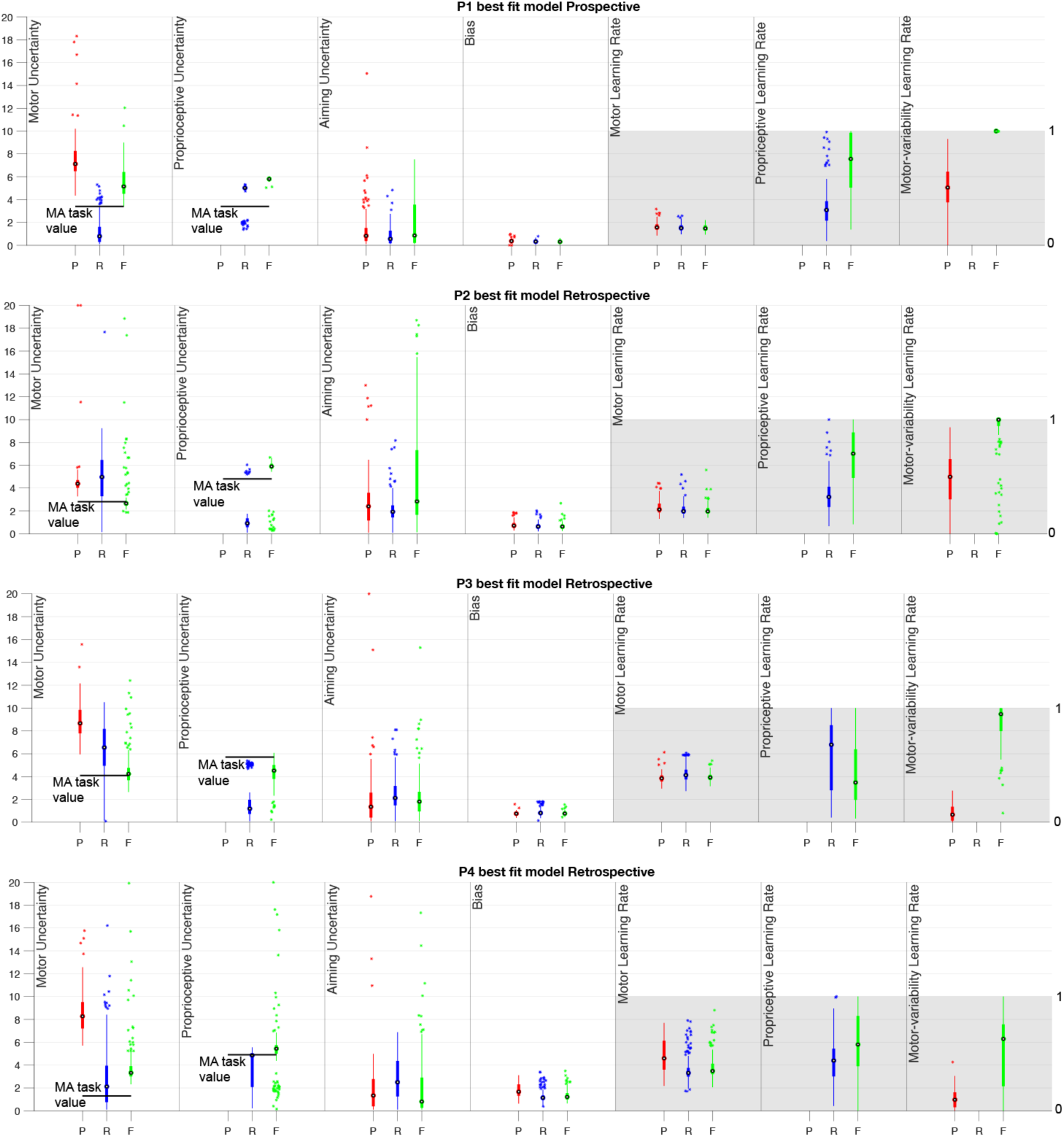
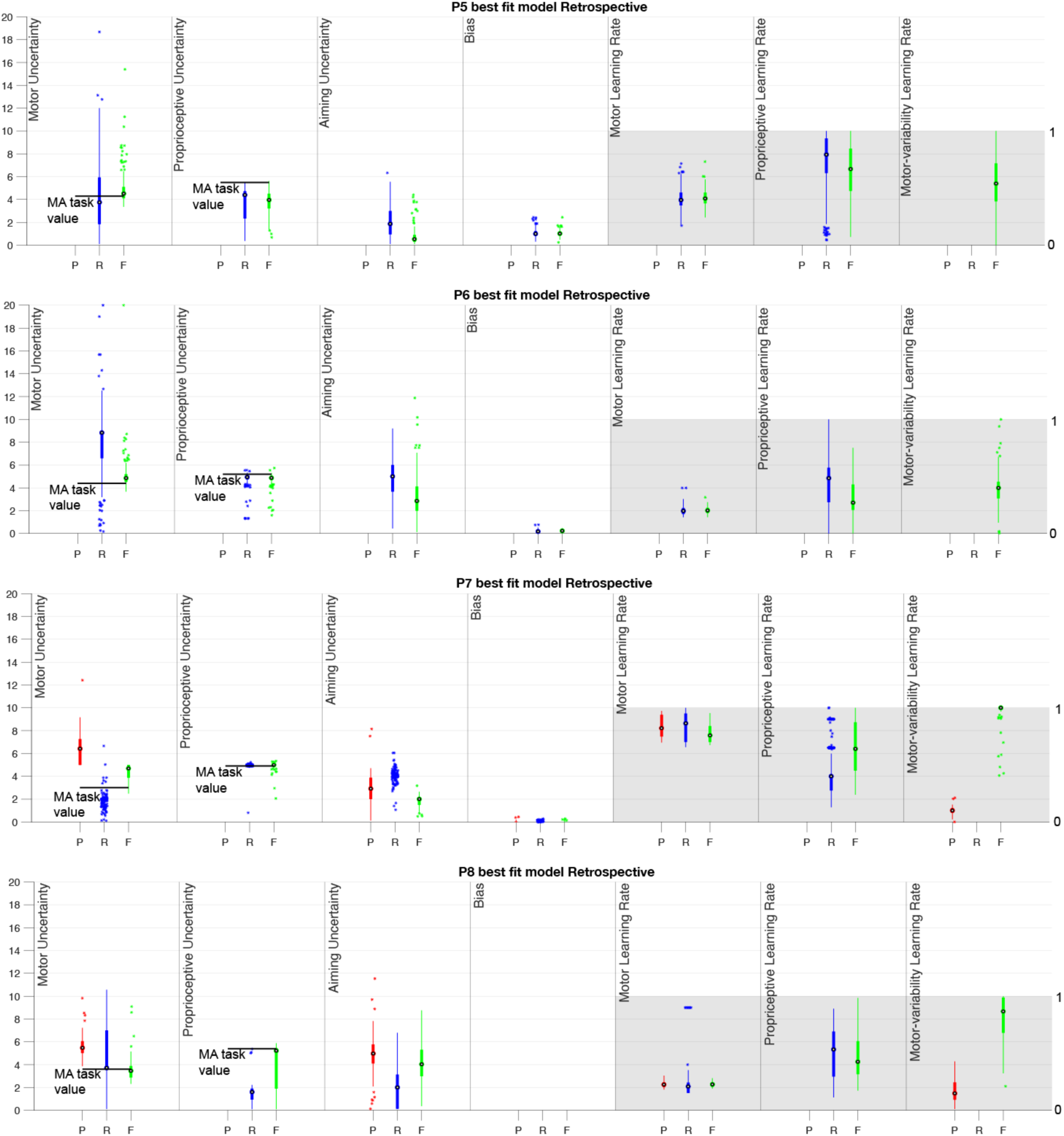
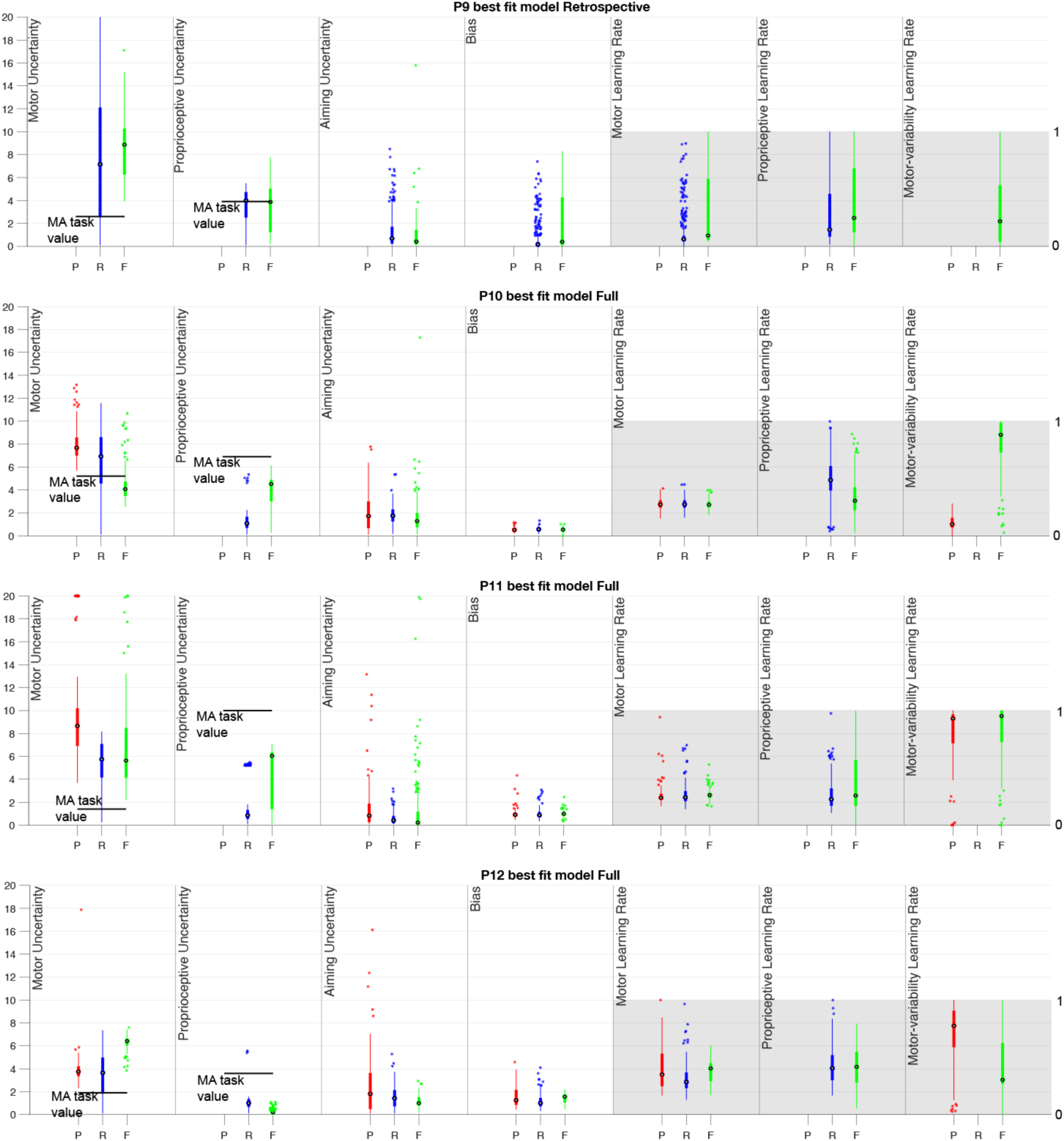
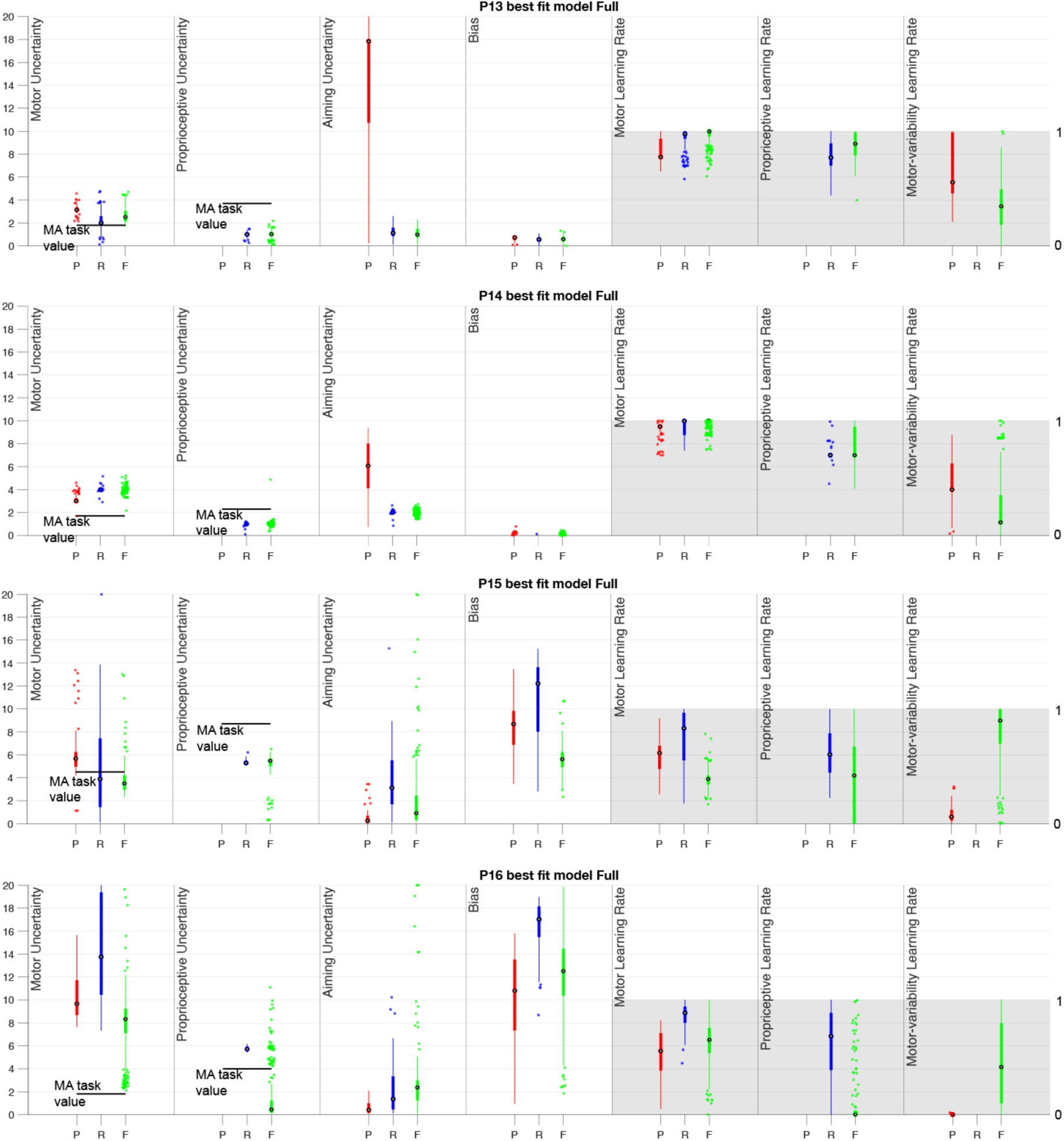
Parameter comparisons for each model fit. For each participant we’ve plotted the range of parameters resulting in a winning iteration of a given model. On the *x*-axis the three models are represented for each parameter, Prospective (P) in red, Retrospective (R) in blue, and Full (F) in green. There are 12 sets of parameters for each iteration so there are between 24 and 444 data points for each model, depending on the number of winning iterations. These box plots show the parameters from the fits that resulted in the winning model producing the lowest least-squares value and winning that round. If the model did not use a particular parameter it has been left blank here. If that participant never had a winning iteration of a given model, there are no winning parameters to show and the model’s section has been left blank. The box edges are the upper and lower quartiles, with the horizontal line reflecting the median, dots representing outliers (calculated using the interquartile range) and whiskers reflecting maximum and minimum values that are not outliers. The black bars in the motor- and proprioceptive-error plots are the values fit for each participant in the separate motor-awareness task. None of the data used to generate those values was used in calculating the parameters shown here and vice versa.

